# The Brain-Derived Neurotrophic Factor Val66Met Genotype Does Not Influence the Grey or White Matter Structures Underlying Recognition Memory

**DOI:** 10.1101/461731

**Authors:** Nicole S. McKay, David. Moreau, Dion T. Henare, Ian J. Kirk

## Abstract

A single nucleotide polymorphism (SNP) in the gene coding for brain-derived neurotrophic factor (BDNF) has previously been associated with a reduction in recognition memory performance. While previous findings have highlighted that this SNP contributes to recognition memory, little is known about its influence on subprocesses of recognition, familiarity and recollection. Previous research has reported reduced hippocampal volume and decreased fractional anisotropy in carriers of the Met allele across a range of white matter tracts, including those networks that may support recognition memory. Here, in a sample of 61 healthy young adults, we used a source memory task to measure accuracy on each recognition subprocess, in order to determine whether the Val66Met SNP (rs6265) influences these equally. Additionally, we compared grey matter volume between these groups for structures that underpin familiarity and recollection separately. Finally, we used probabilistic tractography to reconstruct tracts that subserve each of these two recognition systems. Behaviourally, we found group differences on the familiarity measure, but not on recollection. However, we did not find any group difference on grey- or white-matter structures. Together, these results suggest a functional influence of the Val66Met SNP that is independent of coarse structural changes, and nuance previous research highlighting the relationship between BDNF, brain structure, and behaviour.

## 1. Introduction

Recognition is a critical aspect of memory that allows us to distinguish between items we have experienced before and those we have not. Judgments of recognition are underpinned by two memory subsystems, supporting familiarity and recollection. Familiarity is defined as a type of recognition that allows for the identification of a previously seen stimulus and is often associated with a ‘feeling’ of knowing. In contrast, recollection is a slower, more deliberate process that calls upon associated and detailed information to accompany object identification. For example, recollection could be associated with the specific context the item was last seen in or particular information related to the study period. Familiarity and recollection have been behaviourally and structurally dissociated in previous research (Aggleton & Brown, 1999; Aggleton, Dumont, & Warburton, 2011; Brown & Xiang, 1998; Eichenbaum, Yonelinas, & Ranganath, 2007; Henson, Rugg, Shallice, Josephs, & Dolan, 1999; Mitchell & Johnson, 2009; Wixted, 2007; Yonelinas, 1994; Yonelinas, Kroll, Dobbins, Lazzara, & Knight, 1999; Yonelinas, Otten, Shaw, & Rugg, 2005). For example, it has been suggested that familiarity is an unconscious process dependent upon a diffuse neural network that includes the perirhinal cortex and the uncinate fasciculus, whereas recollection is a conscious and effortful process that relies primarily on structures such as the hippocampus and the cingulate gyrus (Aggleton & Brown, 1999).

Beyond the neural structures supporting recognition memory, a common single nucleotide polymorphism (SNP) in the gene coding for brain-derived neurotrophic factor (BDNF) offers some insight into the genetic basis for variation in recognition memory performance observed across individuals (Egan et al., 2003; Hariri et al., 2003; Toh, Ng, Tan, Tan, & Chan, 2018). This SNP, known as rs6265 or Val66Met, involves a non-synonymous base pair substitution, which results in a change to one of the amino acids of the resulting protein. This switch from a Valine to a Methionine amino acid results in three possible genotypes, Val/Val, Val/Met, and Met/Met. This amino acid substitution is known to result in functional and structural changes in the BDNF protein (Chen et al., 2004). Specifically, it has been reported that this SNP decreases the availability of the mature BDNF protein within the brain (Chen et al., 2005, 2006). This is of particular interest as BDNF is linked to cellular processes that are important for the formation of memory, the maintenance of neuronal connections, and the development of brain structures (Ghosh, Carnahan, & Greenberg, 1994; Hu, Nikolakopoulou, & Cohen-Cory, 2005; Poo, 2001). Given these links, a large body of work has increasingly investigated whether this SNP confers any cognitive disadvantage (Toh et al., 2018).

Previous studies have found that individuals with at least one copy of the Val66Met minor allele (i.e., Val/Met and Met/Met) fare worse on learning and memory tasks (Bekinschtein, Cammarota, Izquierdo, & Medina, 2008; Lamb, Thompson, McKay, Waldie, & Kirk, 2015; Pang & Lu, 2004; Rattiner, Davis, & Ressler, 2005; Spriggs et al., 2019; Tyler, Alonso, Bramham, & Pozzo-Miller, 2002). For example, Egan et al. (2003) reported that Met allele carriers performed significantly worse on episodic memory, as measured by the revised version of the Wechsler Memory Scale (WMS-R). In line with this finding, Hariri et al. (2003) reported a reduction in recognition accuracy and hippocampal activation during a functional-MRI (fMRI) based version of the same task. Further, Ho et al. (2006) found that Met allele carriers have reduced performance on word recognition memory as well as visuospatial tasks. While these findings paint a consistent picture suggesting that the Met allele confers a disadvantage for memory performance, it is important to note that some attempts to replicate these results in clinical populations have not been successful (see Dempster et al., (2005), for example). Furthermore, these studies have measured recognition with a single memory score, despite evidence that recognition is comprised of two distinct memory subprocesses. Given that familiarity and recollection subserve distinct parts of recognition, disentangling the impact of the Val66Met SNP on these processes independently, seems essential. Indeed, it is possible that the influence of the Val66Met SNP is generic across recognition processes, but it could also be that one of these subprocesses drives the overall pattern of reduced accuracy noted in the literature. This is of particular relevance because recollection has been linked explicitly to the hippocampus (Brown & Aggleton, 2001; Yonelinas, 1997), demonstrated in rat studies to be the site of the highest concentrations of BDNF in the brain (Hofer, Pagliusi, Hohn, Leibrock, & Barde, 1990; Maisonpierre et al., 1990).

In line with these behavioural results, structural magnetic resonance imaging (sMRI) studies have provided evidence for differences in grey matter volume between Val/Val individuals and those with one or more copies of the Met allele. Several volumetry studies have reported that Met allele carriers have reduced hippocampal grey matter compared to Val/Val individuals (Bueller et al., 2006; Pezawas et al., 2004; Schofield et al., 2009). Because it is the region in which the highest concentrations of activity-dependent secretion of BDNF are found, the hippocampus is thought to be particularly impacted by Val66Met genotype (Chen et al., 2004). However, the hippocampus is not the only region where Val66Met genotype is reported to influence grey matter volume: Met allele carriers have also been reported to have reduced grey matter volume in the prefrontal cortex (Nemoto et al., 2006), the parahippocampal region (Montag, Weber, Fliessbach, Elger, & Reuter, 2009), as well as a a global reduction in cortical thickness (Yang et al., 2012). Finally, additional evidence to support these findings can be observed in the healthy controls of several clinical studies (Chepenik et al., 2009; Frodl et al., 2007; Ho et al., 2006; Joffe et al., 2009; Matsuo et al., 2009; Szeszko et al., 2005).

The Val66Met SNP is also thought to influence white matter microstructure. Previous research has suggested that BDNF modulates myelinogenesis (Du, Fischer, Lee, Lercher, & Dreyfus, 2003); therefore it is also of interest to consider how this SNP might impact white matter structures within the brain. Several studies have measured diffusion parameters of white matter tracts for Val/Vals and carriers of the Met allele; most commonly, fractional anisotropy (FA) is used as a proxy for tract integrity (Chiang et al., 2011; Forde et al., 2014; Kennedy, Rodrigue, Land, & Raz, 2009; Meng et al., 2017; Montag, Schoene-Bake, Faber, Reuter, & Weber, 2010; Tost et al., 2013; Voineskos et al., 2011). Yet, results have been inconsistent across studies. Some studies have reported an increase in FA (implying higher white matter integrity) in Met allele carriers (Carlson, Cha, Harmon-Jones, Mujica-Parodi, & Hajcak, 2014; Chiang et al., 2011; Tost et al., 2013). For example, Tost et al. (2013) found increased FA in the splenium, posterior thalamic radiation, superior corona radiata, and the superior longitudinal fasciculus. In line with these findings, Chiang et al. (2011) also reported increased FA in the splenium, the superior corona radiata, the left optic radiation and the inferior fronto-occipital fasciculus. In contrast, several studies have reported Met allele carriers to have lower FA in specific white matter tracts such as the uncinate fasciculus (Carballedo et al., 2012), and the anterior corona radiata (Choi et al., 2015), at least in individuals with Major Depressive Disorder. Finally, one study has reported no significant differences in FA between the two genotype groups (Montag et al., 2010).

Given the equivocal results for structural differences between individuals with and without the Val66Met SNP, we sought to quantify structural differences between these groups systematically. We were particularly interested in extending these analyses to regions that underlie recognition memory processing, and in correlating our findings to performance scores on a recognition task. To this end, we used a source memory task to measure both familiarity- and recollection-based recognition memory, and collected sMRI and diffusion-MRI (dMRI) images to obtain structural parameters of the underlying neural circuitry related to each subprocess. We hypothesised that, in line with the dominant previous literature, Val/Val individuals would outperform Met allele carriers on the recognition memory task. Furthermore, we hypothesised that the accuracy difference between our genotype groups would be greatest for recollection judgments than for familiarity judgments, based on the documented link between recollection processes and the hippocampus. Structurally, we hypothesised that Val/Val individuals would show greater grey matter volume, and increased white matter integrity across the brain, and further postulated that this would be particularly evident in the neural circuitry associated with recollection, compared to that of familiarity. In order to test this hypothesis, we included global measures of grey matter volume, together with white matter diffusion parameters–(FA), axial diffusivity (AD), mean diffusivity (MD), radial diffusivity (RD). Each of these diffusion parameters is important for characterising the movement of water within the brain. Previously AD has been linked to axonal damage (Harsan et al., 2006), MD to cellular disarray (Clark et al., 2011), and RD to demyelination (Sun et al., 2006), all important characteristics to consider when using diffusion patterns as a proxy for white matter integrity. We also measured grey matter volume for the hippocampus and parahippocampal region, as these have been directly associated with recollection and familiarity processing (Aggleton & Brown, 1999). Additionally, we measured diffusion parameters across the uncinate fasciculus, cingulate angular bundle, and cingulate gyrus, tracts that have been previously described as linking regions associated with familiarity and recollection processes, respectively (Aggleton & Brown, 1999).

## 2. Methods

### 2.1 Participants

Sixty-one healthy adults, aged 18-33 (*M* = 23.1, *SE* = 0.53), volunteered for this study (see Table 1 for demographics). All participants were recruited at The University of Auckland and reported no history of neurological disorder. Upon genotyping, the participants were divided into two groups, those with the Val/Val genotype, and those with at least one copy of the Met allele. Two participants were unable to complete the behavioural task and were therefore excluded from task-based analyses. All participants gave informed consent, and this study was approved by The University of Auckland Human Participants Ethics Committee.

**Table 1.** General participant information for participants included in the analyses.

|  | Whole sample | Val/Val | Met Carriers | *p*-values |
| --- | --- | --- | --- | --- |
| *N* | 61 | 31 | 30 |  |
| Mean Age (*SE*) | 23.13 (0.53) | 24.03 (0.78) | 22.20 (0.69) | 0.09 |
| Females (Males) | 45 (16) | 21 (10) | 24 (6) | 0.28 |
| Right (Left) Handed | 53 (8) | 26 (5) | 27 (3) | 0.48 |
| Caucasian (Asian) | 53 (8) | 29 (2) | 25 (6) | 0.12 |

### 2.2 Genotyping

Saliva samples were collected from all participants using Oragene-DNA Self Collection kits and stored at room temperature until analysis could take place. Prior to collection, participants were asked not to eat or drink for at least 30 minutes. DNA was extracted from the sample and was resuspended in Tris-EDTA buffer and quantified using a Nanodrop ND-1000 1-position spectrophotometer (Thermo Scientific). Amplification was conducted on the 113bp fragment that coincides with the Val66Met SNP within the BDNF gene. The primers used for this amplification were *BDNF-F 5’-GAG GCT TGC CAT CAT TGG CT-3’ and BDNF-R 5’-CGT GTA CAA GTC TGC GTC CT-3’*. Polymerase chain reaction (PCR) was then applied to the samples, and enzyme digestion was used to cut the samples into the relevant sections. Val fragments of the DNA samples result in two sections of differing lengths, one of 78bps and the other of 35bps, while Met fragments resulted in one section of 113bps in length. DNA was then visualised under ultraviolet light and participants were classified as either Val/Val or Met carriers (those with either Val/Met or Met/Met genotypes). A total of 78 individuals were genotyped for the Val66Met SNP resulting in the following allelic distribution: 43 Val/Val, 29 Val/Met, and 6 Met/Met. This sample was calculated to have a Hardy Weinberg Equilibrium value of 1, and did not deviate from expected distribution values (Petryshen et al., 2010). From this pool, a random subsample of 31 Val/Val homozygotes and 30 Met allele carriers (five homozygotes) were invited to take part in this study.

### 2.3 Source Memory Task

A modified version of the source memory task outlined in Addante et al. (2012) was used. This task requires participants to learn items along with an associated judgment and allows for two types of recognition to be assessed. Item recognition thought to index familiarity, and judgment recognition thought to index recollection. During the study phase of the experiment, participants were presented with 300 objects and were asked to make an associated judgment for each object. On 150 of these trials, participants were asked to decide whether the object presented to them was man-made, and on the remaining 150 trials, participants were asked to decide whether the object presented could fit in a box (size specified by the researcher). Objects were presented to participants in six blocks of 50 stimuli, with breaks of at least 30 seconds between each block. Prior to each stimulus presentation, a fixation cross was presented. The encoding probe (man-made or box) then appeared on the screen for 400 ms. Following this, participants were presented with the object for 1500 ms, and following the trial, participants were asked to respond with a “yes” (press 1) or “no” (press 2) to the cued judgment for that trial. After the study phase, participants were given a 30-minute break with a distraction task (sudoku), and light refreshments were provided. During the retrieval phase, the 300 objects presented in the study phase were mixed with a further 100 novel objects. These 400 objects were presented to the participants in eight blocks of 50. Before each object presentation, a fixation cross was presented for 750 ms. The object was then presented for 1500 ms, and following the end of the trial, participants were asked to decide whether the object presented was from the previous study phase, “old” (press 1), or was a new picture, “new” (press 2). If participants responded that the object was old, a second response screen was presented that asked with which of the two judgments the object was paired within the study phase, “man-made” (press 1), or “box” (press 2). All trials were preceded by a blank screen that lasted between 750 and 1250 ms, to avoid any expectation bias influencing reaction times (see Figure 1).

**Figure 1.**
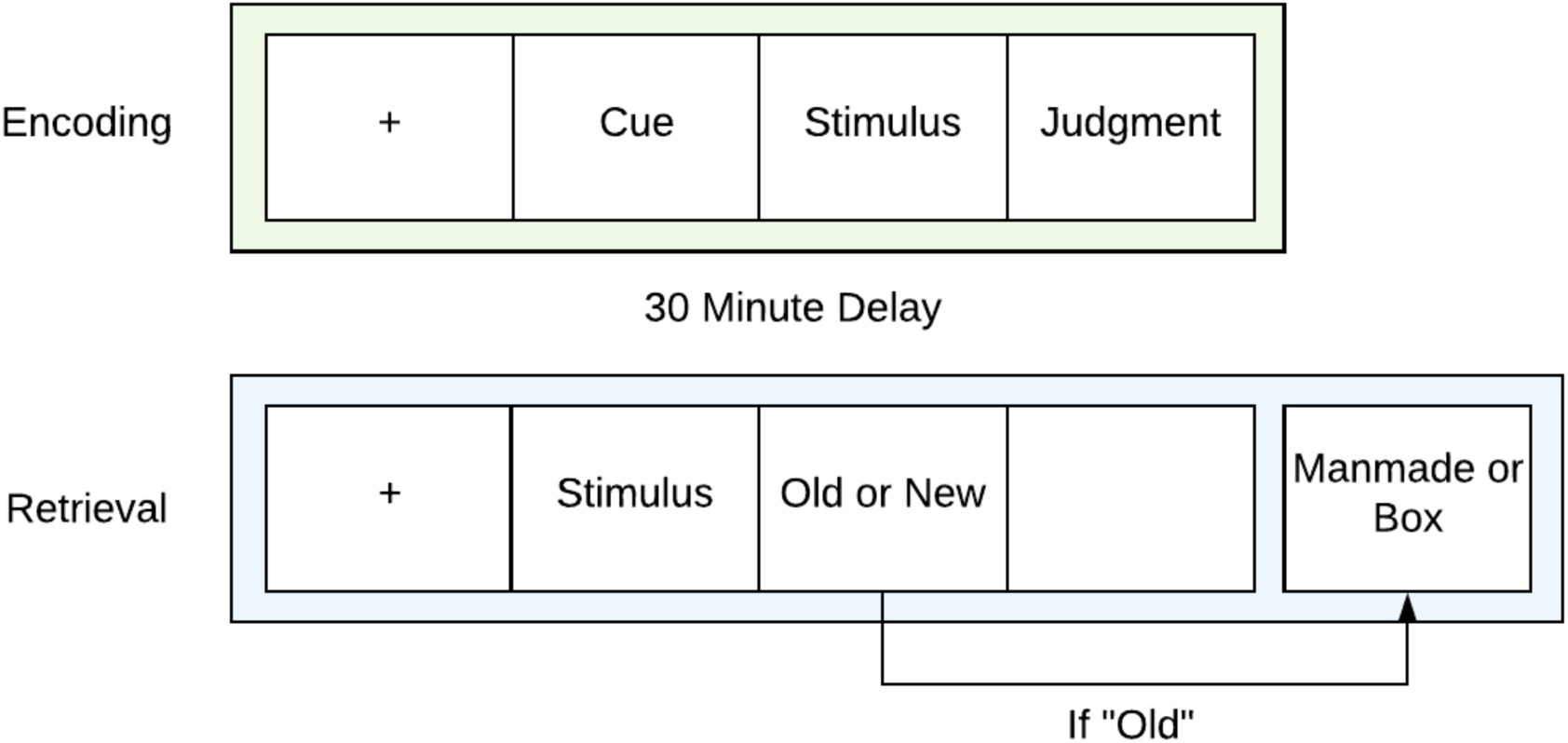
Source recognition memory paradigm. A schematic depiction of the paradigm used for the experiment (adapted from Addante et al., (2012). Participants were shown 300 items in the encoding phase and asked to make an associated judgment for each. After a 30 minute delay, participants then completed the retrieval phase of the experiment.

### 2.4 Stimuli

Picture stimuli were selected from the Bank of Standardized Stimuli (BOSS) (Brodeur, Dionne-Dostie, Montreuil, & Lepage, 2010). This database consists of 930 photos of everyday objects that have been normalised for name, category, familiarity, visual complexity, object agreement, viewpoint agreement, and manipulability (Brodeur, Guérard, & Bouras, 2014). All images are edited so that luminance and colour are equal across the images. Of these 930 photo stimuli, a subset of 500 images was chosen as stimuli for this study. Two images were removed after pilot participants’ feedback that the object was not known in a New Zealand context. Each participant was exposed to a random selection of 400 of these images, 300 in the encoding phase, and a further 100 were introduced in the retrieval phase.

### 2.5 Behavioural Analyses

Performance on the source memory task was used to evaluate differences between the two groups. Mean accuracy was calculated for the old-new judgment, as well as the memory for the associated source judgment, man-made/box. Accuracy on the old-new judgment is thought to index familiarity, while accuracy on the associated source judgment is proposed to index recollection (Addante et al., 2012).

### 2.6 Image Acquisition

The scanning protocol employed for this study is a modified version of the UK BioBank scanning protocol (Miller et al., 2016). Images were acquired on a 3 Tesla Siemens Magnetom Skyra scanner with a 32 channel head coil. Anatomical scans were collected with a T1-weighted magnetised prepared rapidly acquired gradient echo (MPRAGE) sequence. This was collected in a sagittal orientation with the following parameters: TE = 2.83ms, TR = 2000ms, TI = 880ms, FOV = 256mm, Flip angle = 8, slices = 208, GRAPPA acceleration factor = 2, acquisition time = 4m 56s. Diffusion images were collected at an angle approximately parallel to the line joining the anterior and posterior commissures (AC-PC line) using a single shot, simultaneous multi-slice (SMS), spin-echo echo-planar-imaging (EPI) sequence with the following parameters: TE = 92ms, TR = 3600ms, voxel size = 2×2×2mm, FOV = 210×210mm, excite flip angle = 78, refocus flip angle = 160, slices = 72, multiband acceleration factor = 3. The diffusion direction scheme was 100 non-collinear directions at two diffusion weightings (b = 1000, b = 2000 s/mm^2^). Six images were also collected with no diffusion weighting (b = 0 s/mm^2^), three of which had reversed phase encoding to the main imaging sequence, to perform distortion correction.

### 2.7 Preprocessing

All DICOM images were imported to a Linux workstation and converted to NIFTI format using MRIcron (Li, Morgan, Ashburner, Smith, & Rorden, 2016). All subsequent image processing steps were undertaken within FMRIB’s Software Library (FSL) and FreeSurfer (Woolrich et al., 2009). All images were reoriented, and the FSL brain extraction tool (BET) was used to remove the skull and other non-brain structures from the image (Smith, 2002). Each image was checked manually to determine if additional or specialised extraction steps were necessary.

Diffusion images were further processed via the following steps. Eddy current distortions and motion artefacts were corrected for using the FSL Eddy Current tool (Andersson & Sotiropoulos, 2016). Any slices that were detected to have signal loss were replaced by non-parametric predictions generated by Eddy’s underlying Gaussian process. An image with no diffusion weighting for each individual was used to create a binary brain mask (Smith, 2002). FA, MD, AD, and RD maps were generated for each participant using the DTIFIT tool (Behrens et al., 2003). Separately, T1 structural images were also preprocessed using the standard FreeSurfer structural pipeline (FreeSurfer 6.0). Structural images were corrected for any artefacts (Reuter, Rosas, & Fischl, 2010), then a watershed and surface deformation procedure was used to remove the brain from the skull within the image (Ségonne et al., 2004). Images were then registered to Talairach space for subcortical white and grey matter structures to be segmented (Fischl et al., 2002, 2004). A tessellation step was then employed to estimate the grey and white matter structural boundary, and apply any necessary topological correction (Fischl, Liu, & Dale, 2001; Ségonne, Pacheco, & Fischl, 2007). All intersurface boundaries (white matter-grey matter, grey matter-cerebral spinal fluid) were placed in their optimal locations using surface normalisation and intensity gradients (Dale, Fischl, & Sereno, 1999; Dale & Sereno, 1993; Fischl & Dale, 2000). Finally, images underwent surface inflation and registration to a spherical atlas (Fischl, Sereno, & Dale, 1999). Individual participants’ brain tissue segmentation and cortical parcellations were then used as regions of interest for tractography.

### 2.8 Whole-Brain Approaches

#### 2.8.1 Voxel-based morphometry analysis

Voxel-based morphometry (VBM) was conducted using the standard VBM processing pipeline in FSL v 5.0 (Andersson, Jenkinson, & Smith, 2007b; Douaud et al., 2007; Good et al., 2001; Smith et al., 2004). Brain extracted images were segmented into white matter, grey matter and cerebrospinal fluid volume probability maps using FMRIB’s Automated Segmentation Tool (FAST) (Zhang, Brady, & Smith, 2001). A study-specific grey matter template was created using a subset (60) of participants’ images that were chosen from each of the genotype groups. First, these were affinely registered using FMRIB’s Linear Image Registration Tool (FLIRT) into standard space, then concatenated and averaged (Greve & Fischl, 2009; Jenkinson, Bannister, Brady, & Smith, 2002; Jenkinson & Smith, 2001). These images were then re-registered to the new averaged template image using FMRIB’s Non-Linear Registration Tool (FNIRT) (Andersson et al., 2007b). The resulting images were then averaged to create the study-specific template. The grey matter images of all participants were then non-linearly registered to the study template and modulated using the Jacobian of the warp field. The resulting images were concatenated and smoothed, and a general linear model (GLM) approach was used to compare voxel-wise differences in grey matter volume of Val/Val and Met carrier participants, using age as a covariate. Non-parametric statistics were performed using the FSL’s Randomise tool with 5000 permutations and the threshold-free cluster enhancement (TFCE) option (Winkler, Ridgway, Webster, Smith, & Nichols, 2014).

#### 2.8.2 Tract-based spatial statistics

FSL’s tract-based spatial statistics (TBSS) method was used as a whole-brain approach to look at global white matter differences in the preprocessed diffusion images (Jbabdi, Behrens, & Smith, 2010; Smith et al., 2006). Tract-based spatial statistics (TBSS) allows the implementation of a unique nonlinear registration and projection onto an alignment-invariant tract representation (Andersson, Jenkinson, & Smith, 2007a; Andersson et al., 2007b). This technique solves the issue of voxel alignment between participants, ensuring that only voxels that are present in all subjects are included, and does not require smoothing. All FA images were then aligned to the 1×1×1mm FMRIB58_FA standard space target, using a nonlinear registration (Andersson et al., 2007a, 2007b). A standard space version of each subject’s FA image was generated by transforming the FA image to MNI152 standard space (Rueckert et al., 1999). An image containing the mean FA values for all of the subjects was generated and projected onto a mean FA skeleton image, where the skeleton is representative of the centres of all of the tracts common to the subjects. Lastly, for each subject, FA data were projected onto the mean FA skeleton to carry out voxel-wise cross-subject analyses. To also look at how the Val66Met genotype impacts MD, AD, and RD, we used the nonlinear warps and skeleton projection from the FA TBSS analysis to project onto these other diffusion parameter maps.

### 2.9 Specific Region of Interest Analyses

#### 2.9.1 Grey matter volumetric analyses

Region of interest volumetric measures were calculated using Freesurfer 6.0. Of particular interest were grey matter structures within the recognition memory circuits; the thalamus, the hippocampus, the parahippocampal gyrus, the dorsolateral prefrontal cortex, and the entorhinal cortex (see Figure 2). The standard FreeSurfer pipeline for processing structural images was adopted. First images were corrected for any artefacts (Reuter et al., 2010), then a watershed and surface deformation procedure was used to remove the brain from the skull within the image (Ségonne et al., 2004). Images were then registered to Talairach space in order for subcortical white and grey matter structures to be segmented (Fischl et al., 2002, 2004). A tessellation step was then employed to estimate the grey and white matter structural boundary, and apply any necessary topological correction (Fischl et al., 2001; Ségonne et al., 2007). All intersurface boundaries (white matter-grey matter, grey matter-cerebral spinal fluid) were placed in their optimal locations using surface normalisation and intensity gradients (Dale et al., 1999; Dale & Sereno, 1993; Fischl & Dale, 2000). Finally, images underwent surface inflation and registration to a spherical atlas (Fischl et al., 1999). Volumetric measures were collected from these processed images for the regions of interest. The resulting values were analysed using a Bayesian approach to test whether average volume differs between genotype groups.

**Figure 2.**
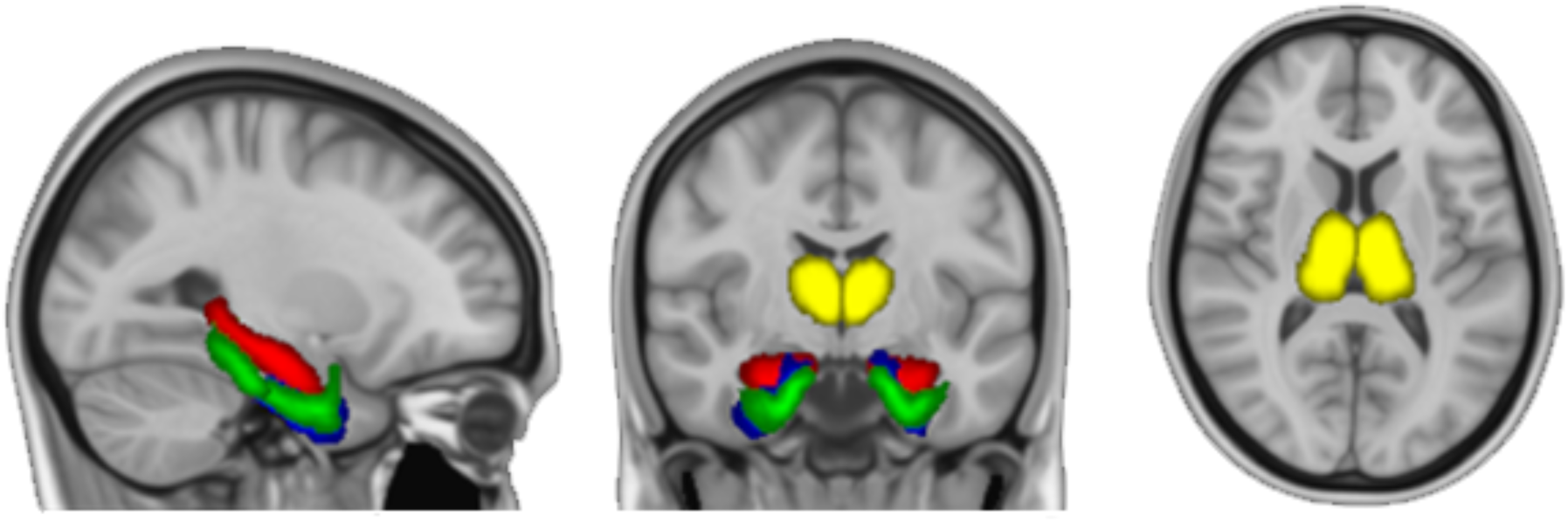
Regions of interest for grey matter analyses. Structures used for the region of interest analyses. In red is the hippocampus, green is the entorhinal cortex, blue is the parahippocampal region (including the perirhinal cortex), and in yellow is the thalamus. In addition to these structures, we also used a section of the dorsolateral prefrontal cortex, which has been excluded from the figure for simplicity.

#### 2.9.2 White matter tractography analyses

White matter pathways associated with memory processing were isolated for region of interest analyses. These included the uncinate fasciculus, cingulum angular bundle, and cingulate gyrus (see Figure 3). Region of interest tractography analyses were conducted using FreeSurfer’s TRActs Constrained by UnderLying Anatomy (TRACULA) tool (Yendiki et al., 2011). TRACULA is an automatic pipeline for reconstruction of major white matter pathways that uses global probabilistic tractography with anatomical priors. These are estimated using FSL’s Bayesian Estimation of Diffusion Parameters Obtained using Sampling Techniques (BEDPOSTX) tool with crossing fibres modelling (Behrens, Berg, Jbabdi, Rushworth, & Woolrich, 2007; Behrens et al., 2003; Jbabdi, Sotiropoulos, Savio, Graña, & Behrens, 2012). An advantage of using TRACULA is that it uses a combination of the prior distributions derived from an anatomical atlas in combination with the cortical parcellation and subcortical segmentation of the individual being analysed. This results in the reconstructed tracts being constrained by the underlying anatomy of the individual.

**Figure 3.**
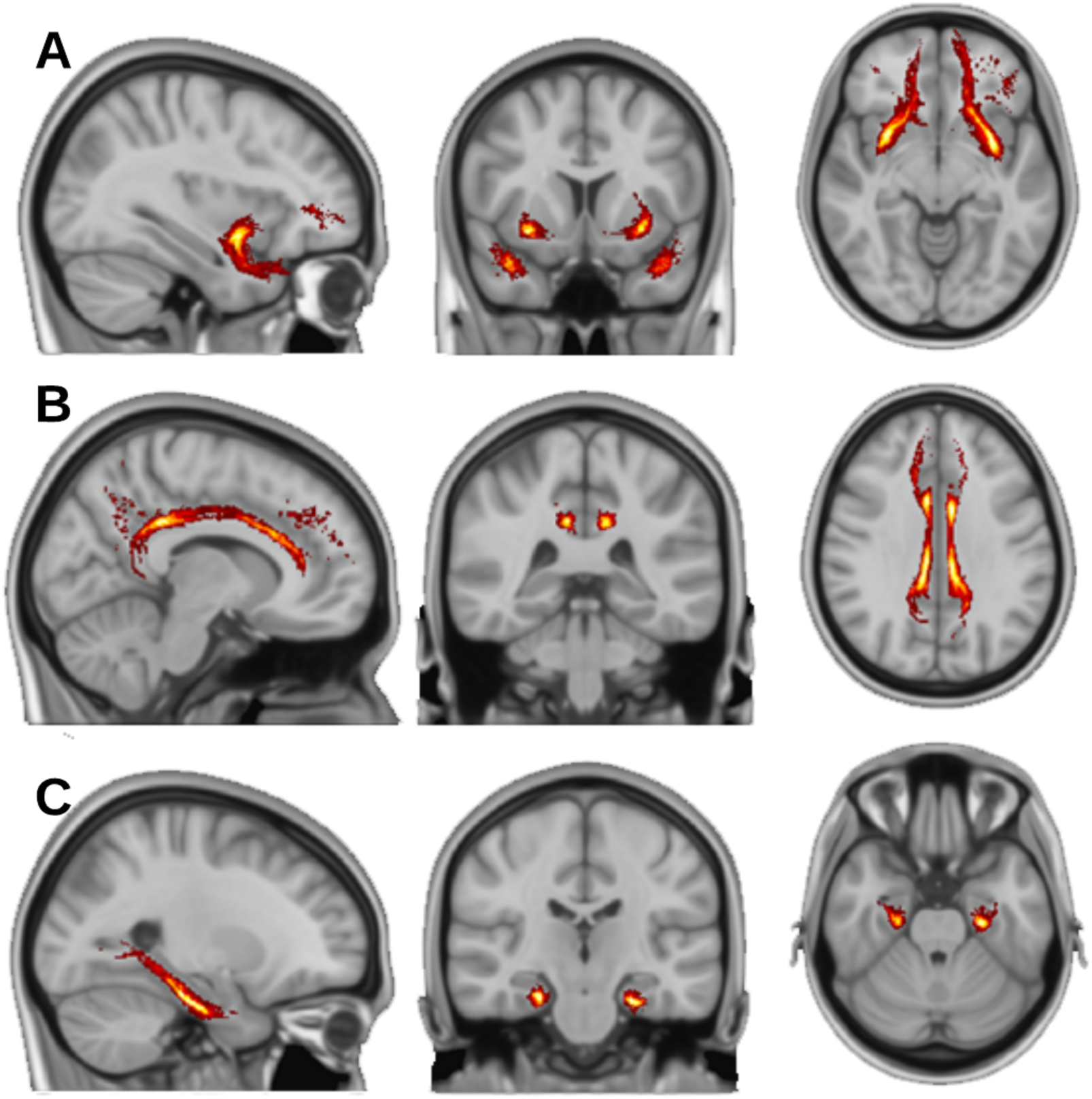
Probabilistic maps of white matter regions of interest. Probabilistic maps depicting the tracts used in the white matter region of interest analyses. The top panel (A) depicts the uncinate fasciculus, the middle panel (B) depicts the cingulate gyrus, and the bottom panel (C) depicts the cingulum angular bundle. The standard MNI152_1mm image is used as a background reference.

### 2.10 Statistics

All statistical analyses were conducted in JASP (JASP Team, 2016), and subsequent plots were made using R (R Core Team, 2018). We used Bayesian statistics for all our analyses to allow quantifying support for the hypotheses we tested–this approach allows a fine-grained representation of the evidence, especially appealing in the cumulative assessment of mixed findings (for a discussion of the advantages of Bayesian analyses over frequentist analyses, see Kruschke, (2013), or Wagenmakers, (2007)). To analyse the behavioural data we used two separate Bayesian independent samples *t*-tests, one for each accuracy measure. To analyse both structural and diffusion data, we used separate Bayesian ANCOVAs, one for each structure (hippocampus, parahippocampal region, thalamus, entorhinal cortex, dorsolateral prefrontal cortex, uncinate fasciculus, cingulate gyrus, and cingulum angular bundle) for each measure of interest (grey matter volume, FA, MD, AD, RD). In the following results section, we report Bayesian model comparisons to quantify evidence for our hypotheses. The resulting Bayes factors are described using the categories proposed by Wagenmakers et al. (2018). Because we understand that some readers may wish to compare these results with the equivalent frequentist analyses, we are providing all of these in the Supplemental Material. In all instances these analyses are broadly consistent with the results presented below.

## 3. Results

### 3.1 Behavioural Analyses

We ran two separate Bayesian independent samples *t*-tests to quantify the evidence for Val66Met group differences in recognition memory. Specifically, we were interested in evaluating the support for the null hypothesis that there is no difference between the two genotypes for our accuracy measures, and also the hypothesis that Val/Val participants scored higher on accuracy measures than Met carriers. For each analysis, the *t*-test was run with the default Cauchy scale (*r* = 0.707; (Morey & Rouder, 2018), a sensible method for estimating the prior distribution when prior values are unknown. For the familiarity measure, we found anecdotal support for a difference in favour of Val/Val participants, whose accuracy scores were higher than Met carriers (BF_10_ = 2.4, *ε* = 6.0×10^-4^%, *M* = 0.82, *SD* = 0.06 and *M* = 0.79, *SD* = 0.09, respectively). For the recollection measure, we found anecdotal evidence for the null hypothesis of no group difference between the two genotype groups (BF_01_ = 1.9, *ε* = 3.0×10^-3^%, *M* = 0.76, *SD* = 0.06 and *M* = 0.74, *SD* = 0.07, respectively). Distributions for the scores of each group for each measure are shown in Figure 4. Posterior and prior distributions, as well as sequential analyses and robustness checks for these tests, are available in the Supplementary Material (Figures SM1 and SM2).

**Figure 4.**
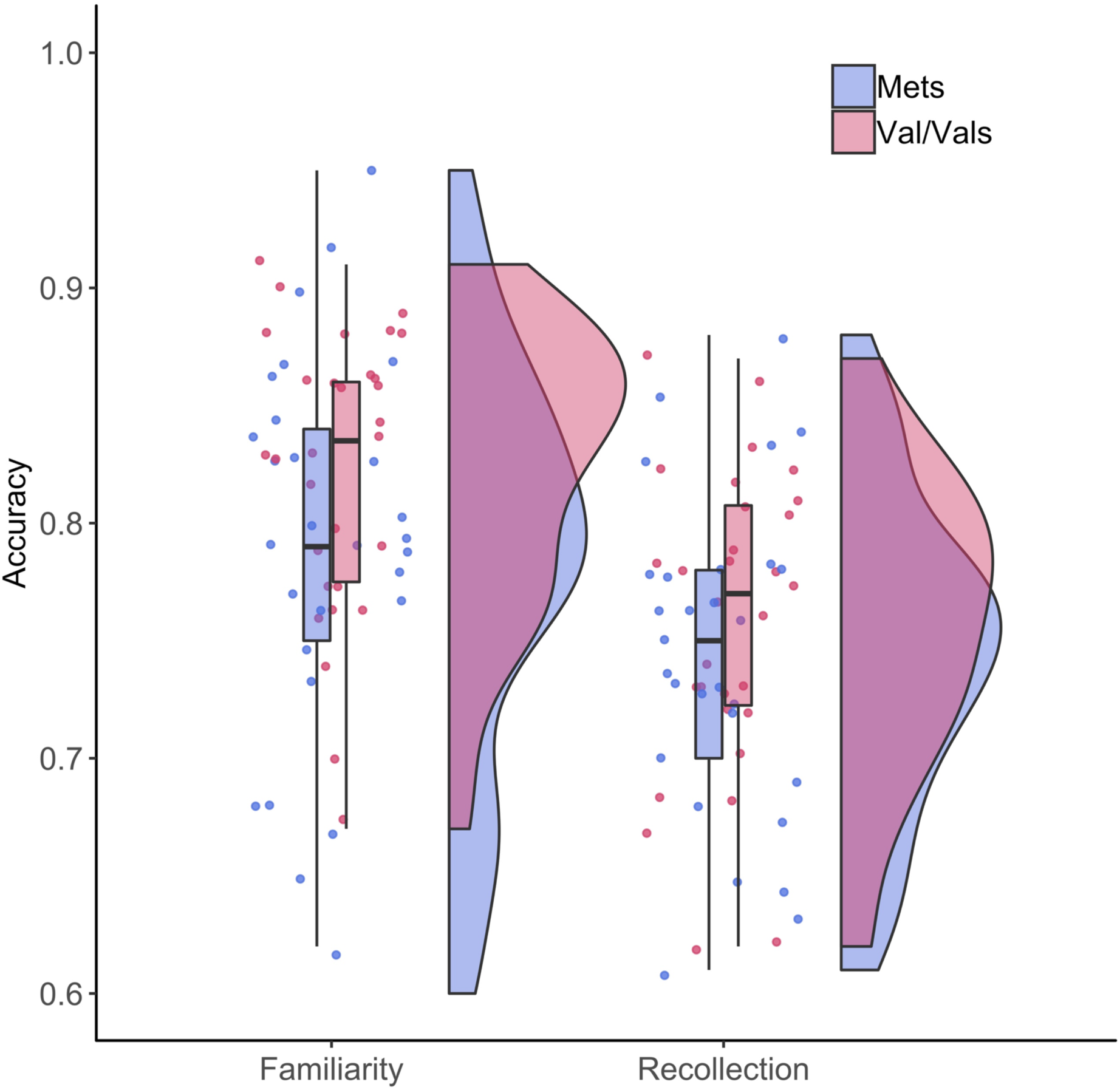
Distributions of behavioural scores. Distributions for the two behavioural measures, familiarity and recollection, split by genotype. Val/Val individuals are shown in red, while Met allele carriers are depicted in blue.

### 3.2 Whole-Brain Analyses

#### 3.2.1 Voxel-based morphometry analysis

A whole-brain grey matter analysis was run to determine if there were any global differences within grey matter regions between participants with the Val/Val genotype and those with at least one copy of the Met allele. Once images were corrected for participants’ ages, and for multiple comparisons, no significant clusters of volumetric differences were found. Clusters that did not survive adjustment for multiple comparisons are presented in the Supplementary Material (Figure SM3).

#### 3.2.2 Tract-based spatial statistics

Whole-brain white matter analyses were run to determine if there were any global differences in the microstructure of white matter pathways between participants with the Val/Val genotype and those with the Met+ genotype. Once images were corrected for participants’ ages, and for multiple comparisons, no significant regions of increased FA were found (Supplementary Material, Figure SM4). To more accurately characterise the properties of the white matter pathways of these groups, MD, RD, and AD maps were also tested for group differences on a global scale. As with the FA analysis, no significantly different voxels were found. Voxels that represented differences that do not survive adjustment for multiple comparisons are presented in the Supplementary Material (Figure SM5).

### 3.3 Region of Interest Analyses

#### 3.3.1 Grey matter volumetric analyses

Structures that are known to be important correlates of recognition memory processing (hippocampus, entorhinal cortex, parahippocampal cortex, thalamus, and dorsolateral prefrontal cortex) were compared for volumetric differences between the two genotype groups (see Figure 5). Here we report the results of five Bayesian ANCOVAs by structure. For each ANCOVA, grey matter volume was used as the dependent variable, genotype (Val/Val, Met+) as the fixed factor, with age and estimated total intracranial volume included as covariates. We used the default prior scales of 0.5 for fixed effects, 1 for random effects, and 0.354 for covariates (Morey & Rouder, 2018). For all regions of interest, we found anecdotal-moderate evidence in favour of the null hypotheses [hippocampus: BF_M_ = 3.43, *p*(M|Data) = 0.77; parahippocampal region: BF_M_ = 2.94, *p*(M|Data) = 0.75; thalamus: BF_M_ = 3.60, *p*(M|Data) = 0.50; entorhinal cortex: BF_M_ = 1.83, *p*(M|Data) = 0.65; DLPFC: BF_M_ = 3.00, *p*(M|Data) = 0.75]. Equivalent analyses controlling for Sex and Ethnicity can be found in the Supplementary Material (Table SM1).

**Figure 5.**
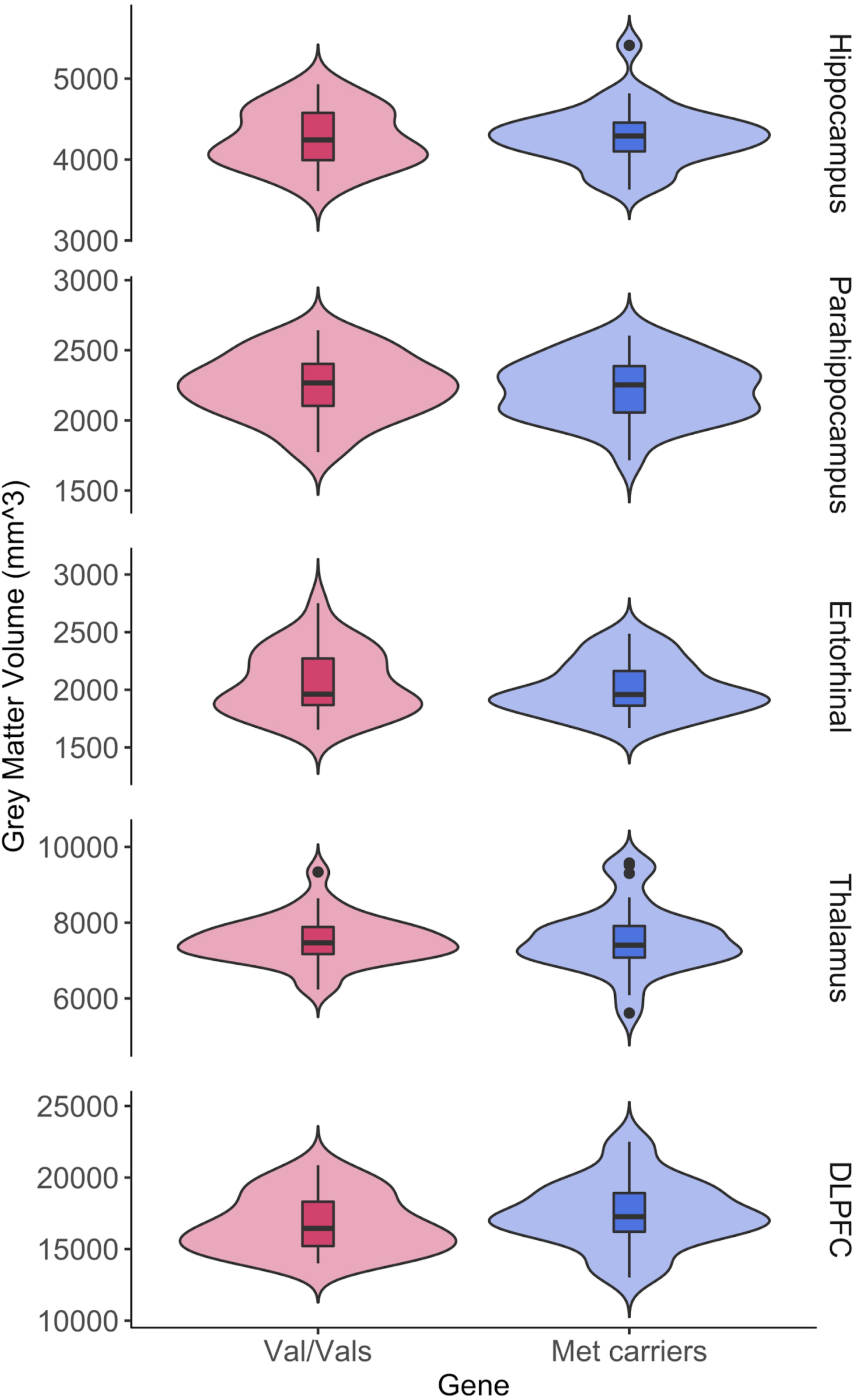
Grey matter volume estimates. Distributions of the grey matter volumes for each genotype (Val/Val vs Met+), across each structure (thalamus, parahippocampal region, hippocampus, entorhinal cortex, and dorsolateral prefrontal cortex). These values split by hemisphere can be viewed in the Supplementary Material (Figure SM6).

#### 3.3.2 White matter tractography analyses

White matter paths that are known to be necessary for recognition processing were reconstructed for ROI analyses to determine if the Val66Met SNP influences white matter tract diffusivity properties. These tracts included the uncinate fasciculus, cingulum angular bundle and cingulate gyrus. For each tract, the average of the tract across both hemispheres was taken for each of the four diffusion parameters. Here we report the results of twelve Bayesian ANCOVAs by tract. For each ANCOVA, the diffusivity measure (FA, AD, MD, RD) was used as the dependent variable, genotype (Val/Val, Met+) as the fixed factor, with age and estimated total intracranial volume included as covariates. We used the default prior scales of 0.5 for fixed effects, 1 for random effects, and 0.354 for covariates (Morey & Rouder, 2018). For all diffusion measures across the uncinate fasciculus, we found anecdotal-moderate evidence in favour of the null hypotheses [FA: BF_M_ = 3.03, *p*(M|Data) = 0.75; MD: BF_M_ = 3.72, *p*(M|Data) = 0.79; AD: BF_M_ = 3.20, *p*(M|Data) = 0.76; RD: BF_M_ = 3.44, *p*(M|Data) = 0.78; see Figure 6]. Similarly, we found support for the null hypothesis across the cingulate gyrus [FA: BF_M_ = 3.46, *p*(M|Data) = 0.78; MD: BF_M_ = 2.84, *p*(M|Data) = 0.74; AD: BF_M_ = 1.71, *p*(M|Data) = 0.63; RD: BF_M_ = 3.82, *p*(M|Data) = 0.79; see Figure 7], and cingulum angular bundle [FA: BF_M_ = 3.31, *p*(M|Data) = 0.77; MD: BF_M_ = 3.78, *p*(M|Data) = 0.79; AD: BF_M_ = 3.62, *p*(M|Data) = 0.78; RD: BF_M_ = 3.61, *p*(M|Data) = 0.78; see Figure 8]. Equivalent analyses controlling for Sex and Ethnicity can be found in the Supplementary Material (Table SM1).

**Figure 6.**
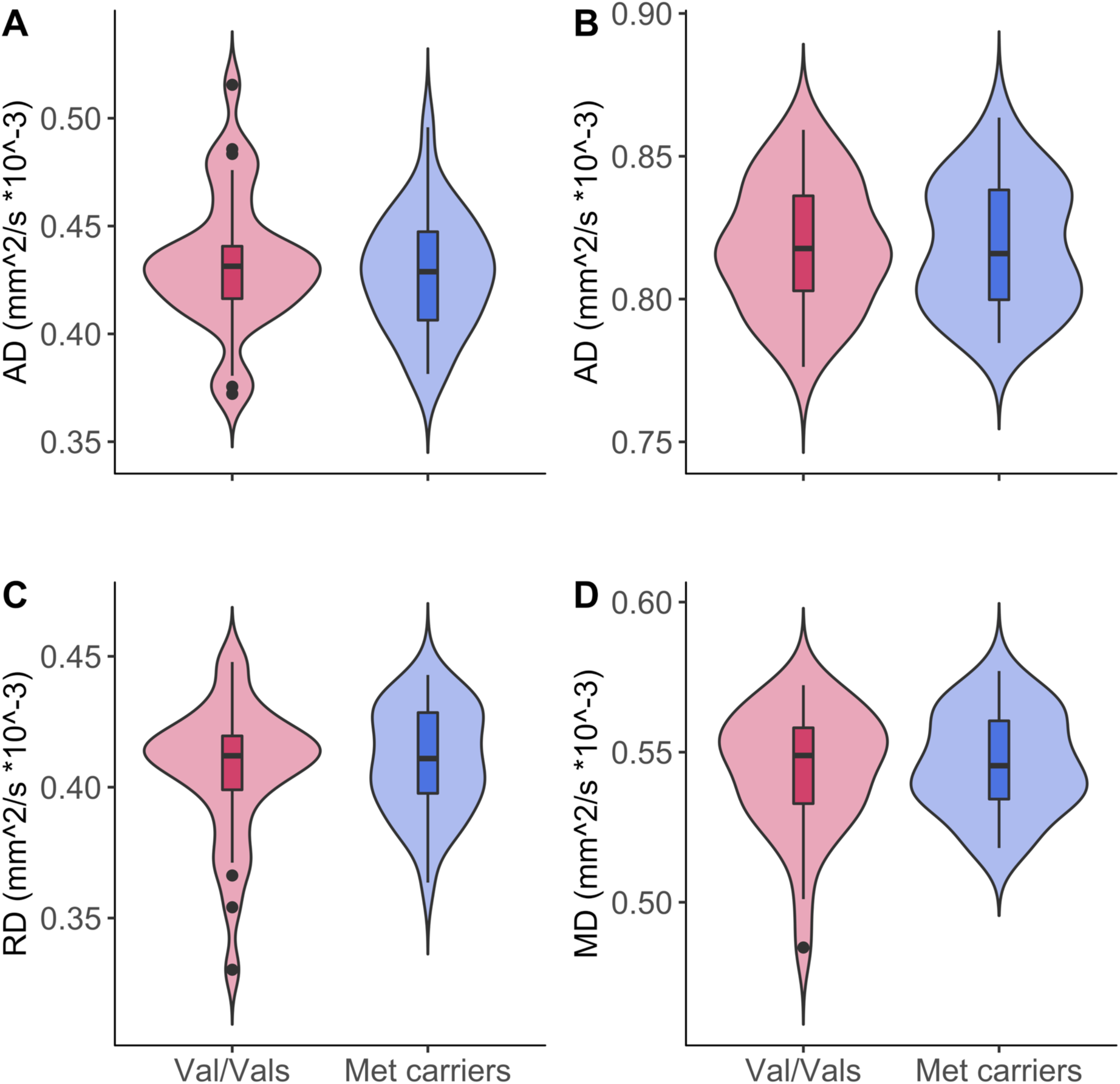
Uncinate fasciculus diffusion estimates. Distributions of the four diffusivity parameters (RD = radial diffusion, AD = axial diffusion, MD = mean diffusion, and FA = fractional anisotropy) measured across the uncinate fasciculus, by genotype. In red are Val/Val participants, and in blue are Met allele carriers. These values split by hemisphere are displayed in the Supplementary Material (Figure SM7).

**Figure 7.**
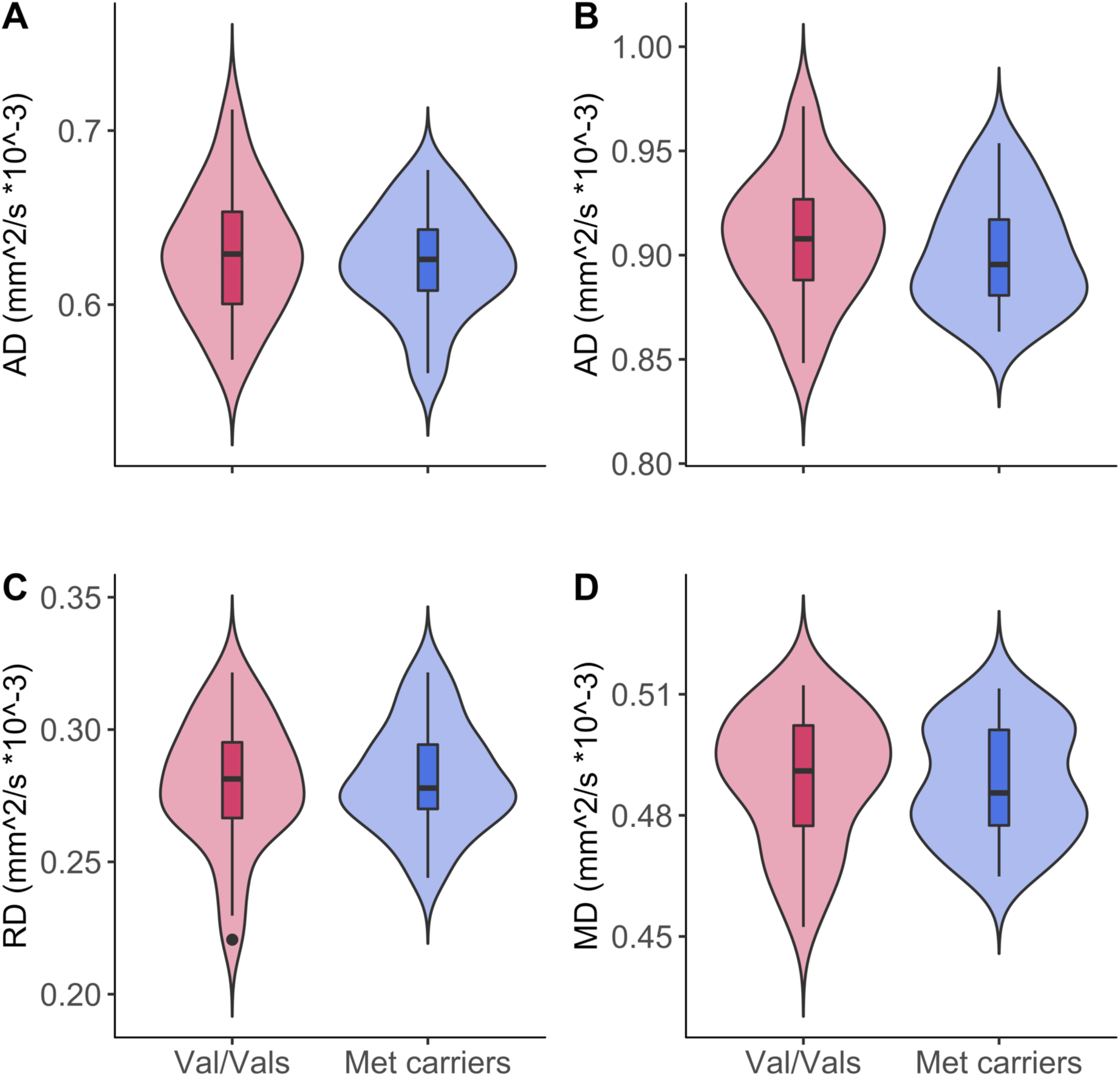
Cingulate gyrus diffusion estimates. Distributions of the four diffusivity parameters (RD = radial diffusion, AD = axial diffusion, MD = mean diffusion, and FA = fractional anisotropy) measured across the cingulate gyrus, by genotype. In red are Val/Val participants, and in blue are Met allele carriers. These values split by hemisphere are displayed in the Supplementary Material (Figure SM8).

**Figure 8.**
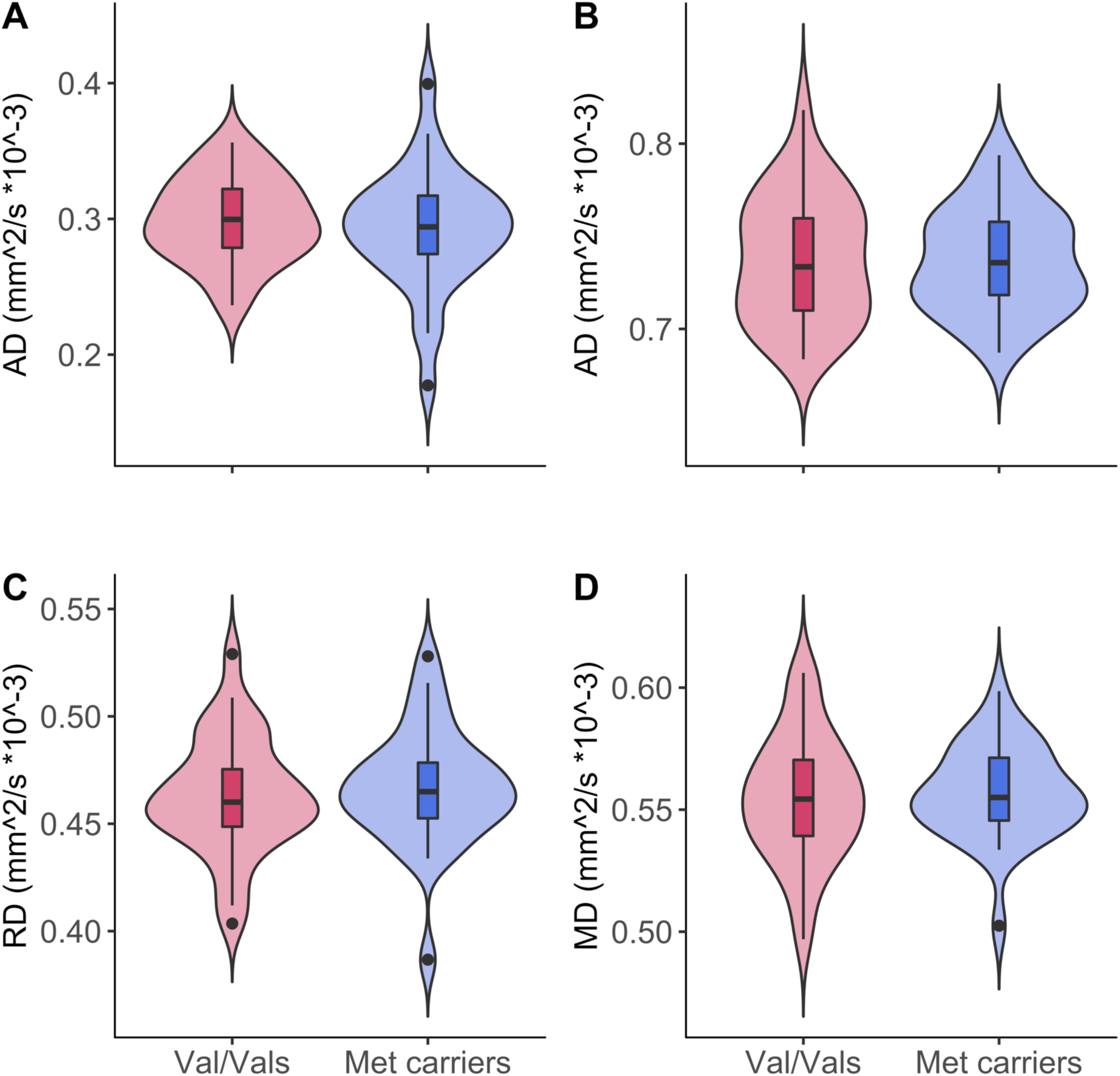
Cingulum angular bundle diffusion estimates. Distributions of the four diffusivity parameters (RD = radial diffusion, AD = axial diffusion, MD = mean diffusion, and FA = fractional anisotropy) measured across the cingulum angular bundle, by genotype. In red are Val/Val participants, and in blue are Met allele carriers. These values split by hemisphere are displayed in the Supplementary Material (Figure SM9).

### 3.4 Behavioural-Structural Correlations

We ran a Bayesian correlation analysis to relate all behavioural scores with average grey matter volume, and diffusivity, for each structure using the default stretched beta prior width of 1 (Morey & Rouder, 2018); see Figure 9 for a summary).

**Figure 9.**
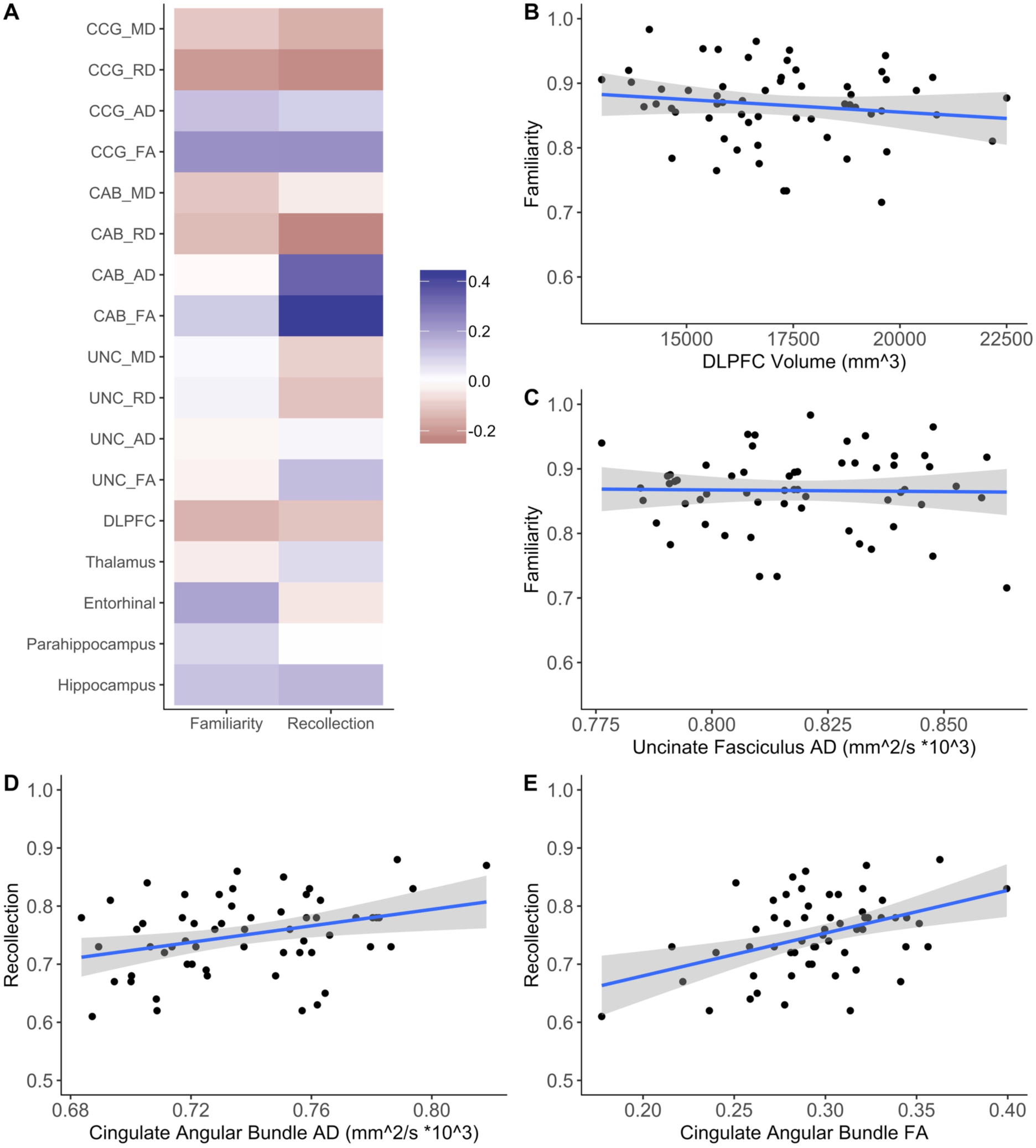
Grey- and white-matter correlation summary. All grey- and white-matter values were correlated with accuracy scores for the familiarity and recollection subtasks. A heat map (panel A) displays the strength of the correlation for all possible combinations. In panels B-E, scatter plots depict all instances where the resulting Bayes Factors show anecdotal-to-strong evidence in favour of the alternative hypothesis. All other correlations showed anecdotal or moderate support for the null hypothesis.

#### 3.4.1 Correlations with grey matter

We found anecdotal evidence of a negative relationship between accuracy on the familiarity task and volume of the DLPFC (*r* = −0.31, BF_10_ = 2.6, 95%CI = −0.54;-0.05), and moderate evidence of no relationship between recollection accuracy and DLPFC volume (*r* = −0.12, BF_01_ = 4.2, 95%CI = −0.35;0.14). All other correlations provided anecdotal-moderate evidence for the null hypothesis of no relationship between accuracy and grey matter volume in the hippocampus (familiarity: *r* = −0.11, BF_01_ = 4.3, 95%CI = −0.35;0.15; recollection: *r* = 0.15, BF_01_ = 3.3, 95%CI = −0.11;0.38), parahippocampal region (familiarity: *r* = −0.16, BF_01_ = 3.1, 95%CI = −0.39;0.10; recollection: *r* = −0.002, BF_01_ = 6.2, 95%CI = −0.25;0.25), entorhinal cortex (familiarity: *r* = −0.08, BF_01_ = 5.2, 95%CI = −0.38;0.18; recollection: *r* = −0.05, BF_01_ = 5.8, 95%CI = −0.29;0.21), and thalamus (familiarity: *r* = −0.17, BF_01_ = 2.8, 95%CI = −0.40;0.09; recollection: *r* = 0.07, BF_01_ = 5.3, 95%CI = −0.18;0.32).

#### 3.4.2 Correlations with white matter

Mean accuracy for both the familiarity and recollection measures were correlated with average scores for each of the diffusion parameters (FA, MD, AD, RD) for each tract (uncinate fasciculus, cingulate gyrus, cingulum angular bundle). We found weak support for a positive relationship between accuracy on the familiarity task and AD within the uncinate fasciculus (*r* = 0.29, BF_10_ = 1.7, 95%CI = 0.03;0.50). Correlations between the two accuracy scores and all other measures within the uncinate fasciculus show anecdotal-moderate support for the null hypothesis (familiarity: FA: *r* = 0.18, BF_01_ = 2.4, 95%CI = −0.07;0.41; RD: *r* = −0.05, BF_01_ = 5.7, 95%CI = −0.30;0.20; MD: *r* = 0.08, BF_01_ = 5.2, 95%CI = −0.18;0.32; recollection: FA: *r* = 0.13, BF_01_ = 3.7, 95%CI = −0.12;0.37; AD: *r* = 0.02, BF_01_ = 6.1, 95%CI = −0.23;0.27; RD: *r* = −0.12, BF_01_ = 4.3, 95%CI = −0.35;0.14; MD: *r* = −0.09, BF_01_ = 5.0, 95%CI = −0.33;0.17). When correlating the diffusion parameter values of the cingulum angular bundle to accuracy on the two recognition tasks we found moderate support for a positive relationship between AD and recollection accuracy (AD: *r* = 0.33, BF_10_ = 4.2, 95%CI = 0.08;0.53), and strong support for a positive correlation between FA and recollection accuracy (FA: *r* = 0.43, BF_10_ = 42.8, 95%CI = 0.19;0.61). All other correlations with measures of diffusion within this tract showed weak-moderate support for the null hypothesis of no relationship between recognition scores and diffusivity (familiarity: FA: *r* = −0.07, BF_01_ = 5.4, 95%CI = −0.31;0.19; AD: *r* = 0.20, BF_01_ = 2.1, 95%CI = −0.06;0.42; RD: *r* = 0.20, BF_01_ = 2.0, 95%CI = −0.06;0.42; MD: *r* = 0.25, BF_01_ = 1.0, 95%CI = −0.01;0.47; recollection: RD: *r* = −0.23, BF_01_ = 1.3, 95%CI = −0.45;0.02; MD: *r* = −0.04, BF_01_ = 5.9, 95%CI = −0.28;0.22). Finally, all correlations between between diffusion within the cingulate gyrus and familiarity or recollection accuracy showed weak-moderate support for the null hypothesis of no relationship (familiarity: FA: *r* = 0.12, BF_01_ = 4.2, 95%CI = −0.14;0.36; AD: *r* = 0.16, BF_01_ = 3.0, 95%CI = −0.10;0.39; RD: *r* = −0.09, BF_01_ = 4.9, 95%CI = −0.33;0.17; MD: *r* = 0.02, BF_01_ = 6.1, 95%CI = −0.24;0.27; recollection: FA: *r* = 0.23, BF_01_ = 1.3, 95%CI = −0.03;0.45; AD: *r* = 0.10, BF_01_ = 4.7, 95%CI = −0.16;0.34; RD: *r* = −0.22, BF_01_ = 1.5, 95%CI = −0.44;0.04; MD: *r* = −0.15, BF_01_ = 3.2, 95%CI = −0.38;0.11).

## 4. Discussion

This study aimed to replicate three major findings in the BDNF and memory literature. Firstly, that carriers of the Met allele have decreased performance on recognition memory tasks, secondly, that they show reduced grey matter volume in structures that are thought to underpin memory, and finally, that they have reduced white matter integrity. We aimed to further complement these previous findings by considering the familiarity and recollection aspects of recognition memory separately. Our reasoning for this was based on evidence that recollection is dependent upon the hippocampus, a region known to have the highest level of activity-dependent BDNF secretion (Chen et al., 2004; Egan et al., 2003), while familiarity is more dependent on extra-hippocampal structures such as perirhinal cortex (Aggleton & Brown, 1999). Given that the Met allele is known to reduce secretion, impair folding, and slow the trafficking of the BDNF protein, it was thought that the hippocampus and the processes it subserves would be most impacted by the cellular changes this SNP confers. It was therefore of interest to explore whether Val66Met genotype might have a greater influence on recollection-compared to familiarity-based recognition responses. We used a source recognition memory task to measure familiarity and recollection accuracy for each individual. Whole-brain grey matter volumes and white matter diffusion parameters were calculated for global comparisons. Grey matter volumes were also calculated for the left- and right-thalamus, hippocampus, parahippocampal region, and entorhinal and dorsolateral prefrontal cortices. These are all structures highlighted in neuroanatomical accounts of recognition memory (Aggleton & Brown, 1999). To examine how this genotype influences white matter integrity in specific paths underlying recognition memory, we measured diffusivity parameters across pathways that connect regions that are critical for memory processing. Specifically, we reconstructed the uncinate fasciculus, a tract connecting the medial temporal lobe to the prefrontal cortex, and cingulum angular bundle and cingulate gyrus, two tracts that are responsible for linking the prefrontal cortex to the hippocampus via the anterior thalamus (estimates for these structures, split by hemisphere, are displayed in Figures SM16:18 within the Supplementary Material).

Behaviourally, we replicated the finding that Met allele carriers have lower accuracy on recognition tasks. Interestingly, we found this to be true for our familiarity measure, but not our measure of recollection. Initially, we had hypothesised that when recognition was broken into its two subcomponents, Met allele carriers would show the greatest deficits on recollection memory judgments. As noted above this hypothesis was based on evidence that the Met allele is associated with lower levels of the BDNF protein during activity in the hippocampus (Chen et al., 2004), a region specifically linked to recollection. Furthermore, given that recollection reflects deeper, more complex, processing, we hypothesised that this measure would be more sensitive to detect small differences in accuracy between groups. However, our results suggest that the Val66Met genotype has a stronger association with the familiarity subtask, which required the identification of a previously seen object. This indicates that once an item is recognised as either old or new, both genotype groups are equally able to recall associated memory details, and suggests that Met allele carriers have more difficulty recalling weak memory traces. This is consistent with previous research using the source memory task, which has described weaker memory traces as those items that are recognised as old without any associated source memory (Addante et al., 2012). Given that we used the item identification and source recognition judgments as respective proxies for familiarity and recollection, our results are consistent with this prediction.

Previous research suggested that volumetric differences related to Val66Met genotype could be detected in grey matter structures (Bueller et al., 2006; Montag et al., 2009; Nemoto et al., 2006; Pezawas et al., 2004; Schofield et al., 2009; Yang et al., 2012). In the present study, we used a VBM approach to assess global grey matter characteristics for each group; however, we did not find any regions of significant volumetric differences between Val/Vals and Met carriers. In order to look at familiarity and recollection subcomponents, we also took a region of interest approach, which allowed us to assess grey matter volume in memory-related structures. Five structures—the thalamus, hippocampus, parahippocampal region, entorhinal cortex and dorsolateral prefrontal cortex—were chosen based on Aggleton and Brown’s (1999) neuroanatomical account of the dual process model of recognition. In all analyses, we did not find evidence to support the previous reports that Met allele carriers have reduced grey matter volume compared to Val/Val individuals. Of particular interest is our failure to replicate the previous finding that Val66Met genotype impacts hippocampal volume (Bueller et al., 2006; Pezawas et al., 2004; Schofield et al., 2009). Although hippocampal volume differences are often cited in the BDNF literature as one of the most established effects of the Val66Met SNP, our findings are nonetheless in line with several studies that have failed to replicate this result (Harrisberger et al., 2014; Karnik, Wang, Barch, Morris, & Csernansky, 2010; Richter-Schmidinger et al., 2011). Furthermore, several clinical studies have also failed to replicate this result within their control participants (Benjamin et al., 2010; Dutt et al., 2009; Gruber et al., 2012; Jessen et al., 2009; Koolschijn et al., 2010). These results may suggest that the link between hippocampal volume and the Val66Met SNP is not as reliable as previously thought.

Val66Met genotype has also previously been shown to influence the integrity of white matter tracts (Carlson et al., 2014; Chiang et al., 2011; Forde et al., 2014; Meng et al., 2017; Tost et al., 2013). In order to further corroborate this finding, we first used a TBSS approach to assess global white matter diffusivity for each group. We looked at each of the four main diffusion parameters; FA, MD, RD, and AD. Using this whole-brain approach, we did not find any regions of significantly different diffusion patterns between our two groups. We also used probabilistic tractography to reconstruct the uncinate fasciculus, cingulum angular bundle, and cingulate gyrus for each participant, given the reports linking these tracts with recognition processing. The uncinate fasciculus is a white matter tract that connects the prefrontal cortex to the perirhinal cortex; areas that are associated with familiarity and item recognition processing (Aggleton & Brown, 1999). Both the cingulate gyrus and cingulum angular bundle are white matter tracts that are known to connect the prefrontal cortex to the hippocampus via the anterior thalamus; regions that are specifically linked to recollection. As with our grey matter regions of interest, our choice of tracts is based upon the neuroanatomical account of recognition proposed by Aggleton & Brown (1999). In line with the whole-brain approach, we characterised water movement within these tracts using four parameters; FA, MD, AD and RD. We found no differences in the diffusion parameters of our two groups across these tracts. This contradicts previous work that has reported an increase in FA for Met allele carriers across the uncinate fasciculus (Carlson et al., 2014). However, our results do support other reports showing the absence of differences between these two gene groups (Montag et al., 2010). Using a Bayesian approach, we were able to circumvent many of the typical limitations associated with the assessment of null findings, to instead quantify the evidence for the absence of group differences. In all instances, we found varying degrees of evidence for the null model that there is no Val66Met genotype impact on grey matter volume or white matter diffusivity. Together with the outcome of our whole brain analyses, we take these regions of interest results as evidence that within our sample, Val66Met genotype does not appear to influence the volume or connectivity of brain structures.

One interesting implication of our results is that they provide support for reduced recognition memory in Met allele carriers in the absence of any detectable structural differences. This implies that the mechanism driving the behavioural difference noted in these individuals is not dependent on large-scale grey matter volumetric differences or reduced white matter integrity. This directly contradicts previous studies that have suggested that the role of the BDNF protein in the development and maintenance of global brain structures could be a potential mechanism underpinning the behavioural differences observed in carriers of the Met allele. Furthermore, previous researchers have postulated that the impact of this SNP should be greatest at the neurophysiological level given this is less removed from the biological impact of gene transcription, compared to processes of a higher level such as cognition (Hariri, Drabant, & Weinberger, 2006). This idea has previously been used as support for research proposing that structural deficits would underpin the behavioural differences observed in these groups (Kambeitz et al., 2012). However, in line with this, and given that the Met allele has been shown to impair the functionality of the BDNF protein as well as it’s concentration within memory-related structures, we propose it is possible that the behavioural effects that are being measured are the result of a short-term, functional, impact of this allele. One such mechanism that might underpin our behavioural difference could be related to the relatively low level of BDNF protein available in Met allele carriers. Importantly, this is not something that can be directly explored with the current data.

There are several limitations that should be acknowledged. First, the task we used aimed to separate familiarity and recollection response types by taking accuracy from the item recognition judgment as a measure of familiarity, and accuracy from the associated judgment as a measure of recollection. However, it is unlikely that familiarity and recollection are completely distinct in this task as they are performed in a serial manner. That is, for each object, participants are first asked whether they recognise an object, before being asked to identify the associated judgment that was made during encoding. A further issue with this task is that it may be better at detecting recollection dependent responses than familiarity responses. The encoding phase of this task requires participants to mentally manipulate the items in a way that could bias them to recall associated information about the item at the first instance of recognition. That is, participants may be unable to prevent recall of the associated encoding context when the item is presented for the first recognition task. We attempted to mitigate the impact of these two limitations by using a measure of familiarity that is thought to be free of the influence of strong recollection (Addante et al., 2012). Our familiarity score was derived from trials that included correct item identification in the absence of a correct source recognition judgment. These trials are thought to index familiarity without deeper recollective processing. Further, our recollection measure was derived from trials where both item recognition and source recognition judgments were correctly reported. While this means that our recollection score cannot speak to recollection in the absence of familiarity, as some previous research has (Addante et al., 2012), it is commonly accepted that familiarity is present in recollection responses.

A second limitation is that we grouped our participants into Val/Vals and carriers of the Met allele. While it would be of great interest to split participants into the three possible genotypes and explore the potential dose-like effect of the Met allele, it was not possible with our current sample size. However, in support of our research design, many previous studies have also split their data in this way (Bueller et al., 2006; Frodl et al., 2007; Szeszko et al., 2005). Furthermore, studies that did investigate the possible dose effect of the Met allele have provided inconsistent results (Beste, Baune, Domschke, Falkenstein, & Konrad, 2010; Chen et al., 2006; Egan et al., 2003; Forde et al., 2014). Given the relatively low frequency of Met/Met individuals (~1-4% of the population; (Mukherjee et al., 2011)), this inconsistency across studies could be due to extremely heterogeneous samples. Despite this, of particular relevance are the findings of Forde et al. (2014), that reported no dose effect for the Val66Met SNP. Forde et al. (2014) observed that Val/Met individuals have the most distinct grey- and white-matter structures compared to either Val/Val or Met/Met individuals. Contrary to previous studies that investigated the dose effect of the Met allele, Forde et al. (2014) was the first study to recruit balanced genetic groups. Importantly, this allowed for the first unbiased analysis of whether an allele dose effect exists for the Val66Met SNP. While the results of the Forde et al. study suggest there is no dose effect for the Met allele, it does also raise questions about the relationship between having one and two copies of the Met allele that needs further addressing, and is beyond the scope of the current experimental design.

Finally, it is important to acknowledge our relatively low sample size. Given previous meta-analyses on this topic have shown consistently small effect sizes (Hajek, Kopecek, & Höschl, 2012; Harrisberger et al., 2014; Kambeitz et al., 2012; Molendijk et al., 2012), it is clear that the current study design would have benefitted from a larger sample size. Despite this, a major strength of using Bayesian analyses is that sample size is factored into the calculation of Bayes factors–with low sample size, Bayes factors tend to favour neither the null nor the alternative hypothesis. Our reported Bayes factors provided anecdotal-moderate support for the null hypotheses, and therefore instil some confidence that our study design is sensitive enough to be interpreted in a meaningful way. Furthermore, we argue that by providing Bayes factors we enable cumulative assessments of evidence that can take into account positive and null findings. Given this, we urge our results should not be considered in isolation, but instead as part of the wider body of literature on this topic.

There is some controversy in the BDNF literature as to what influence the Val66Met SNP has on the grey- and white-matter structures within the brain. In an attempt to help resolve this, our study aimed to replicate three major findings, that those carrying at least one copy of the Val66Met minor allele have lower accuracy on recognition memory tasks, reduced grey matter volume, and reduced white matter integrity. We set out to further these results by also breaking down the recognition measures into familiarity and recollection specific measures, as well as measuring grey- and white-matter measures for structures critical to these behaviours separately. Our results replicated the behavioural findings of previous researchers, with Met allele carriers having lower accuracy, specifically in the familiarity subset of the task. However, contrary to many previous findings, we did not observe any regions of structural differences for this group, suggesting that the impact of the Val66Met SNP on memory might be best characterised as an acute influence, possibly linked to the differential availability of BDNF impacting synaptic plasticity during memory formation. It would be of interest for future extensions of this research to include additional measures such as from fMRI or EEG, in order to directly explore the functional impact of the Val66Met SNP during recognition. Furthermore, while our structures were selected based on a neuroanatomical account of recognition (Aggleton & Brown, 1999), it is possible an fMRI study could identify other structures that are important for recognition memory processing, and therefore would be of interest to investigate too.

## Supporting information

Supplemental Analyses

## Acknowledgements

The authors would like to thank all of the participants who took part in this study, as well as the staff at the Centre for Advanced MRI (CAMRI) for their support throughout this project.

## Data availability statement

Due to restrictions imposed by the University of Auckland Human Participants Ethics Committee, participant-level genetic and MRI data are not available for sharing. However, aggregate data and research code is available upon request, by contacting the corresponding author.

