## Supplemental Analyses for "The Brain-Derived Neurotrophic Factor Val66Met Genotype Does Not Influence the Grey or White Matter Structures Underlying Recognition Memory"

### Supplementary Material

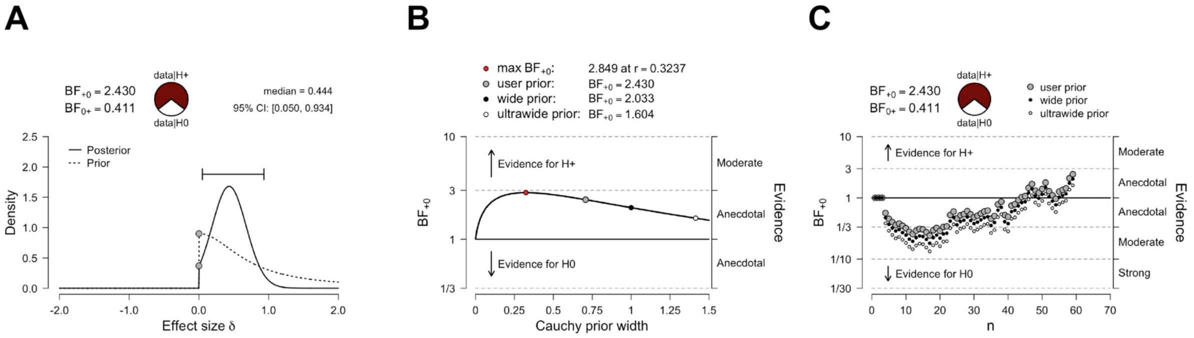


**Figure SM1**. **Supporting plots for the familiarity *t*-test**. Panel A: Prior and posterior distributions for the *t*-test conducted on the familiarity scores. Panel B: Robustness checks showing how the resulting Bayes factor changes across a wide range of priors. Panel C: A sequential analysis depicting the evidence for each of the null and alternative hypotheses, as sample size increases. Accompanying Bayes factors and the credible interval are also displayed.

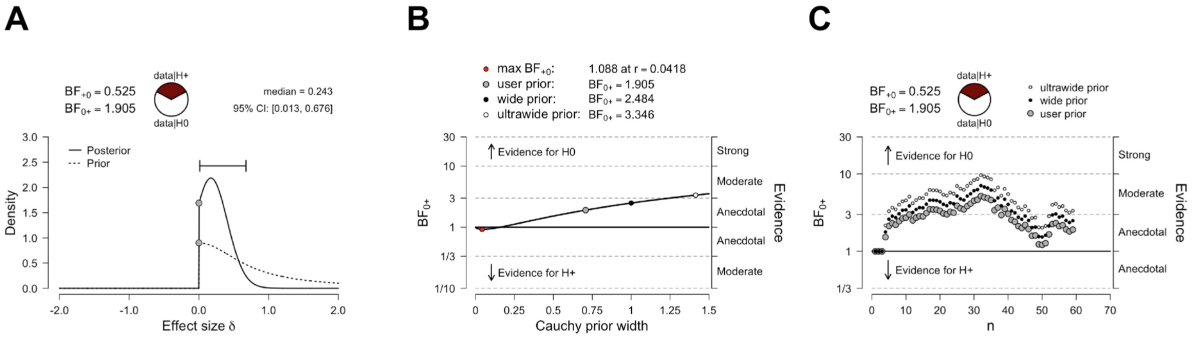


**Figure SM2**. **Supporting plots for the recollection *t*-test**. Panel A: Prior and posterior distributions for the *t*-test conducted on the recollection scores. Panel B: Robustness checks showing how the resulting Bayes factor changes across a wide range of priors. Panel C: A sequential analysis depicting the evidence for each of the null and alternative hypotheses, as sample size increases. Accompanying Bayes factors and the credible interval are also displayed.

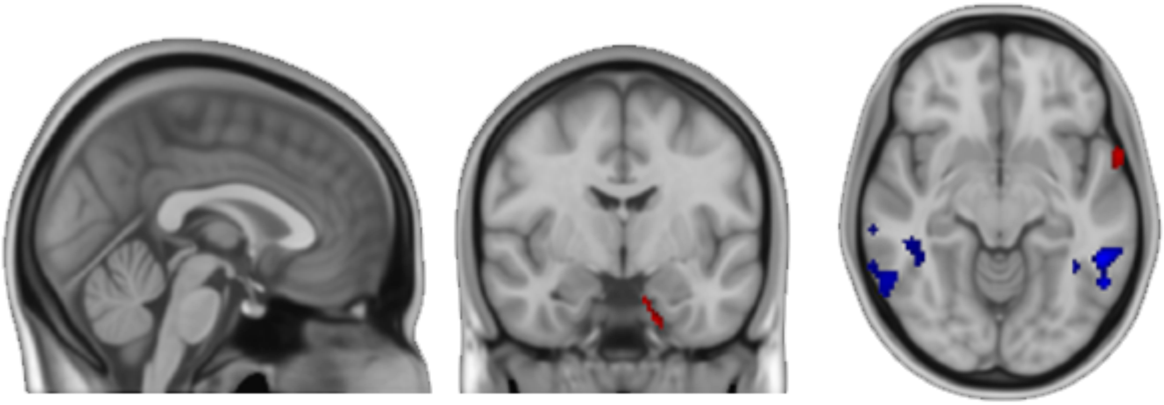

**Figure SM3. VBM results prior to multiple comparison correction**. Depicted are the results of the VBM analysis prior to multiple comparison corrections. The cluster in red indicates greater volume for Val/Val participants compared to Met carriers; clusters in blue represent areas where Met carriers have greater volume compared to Val/Val participants. The standard MNI152_1mm image is used as the background template.

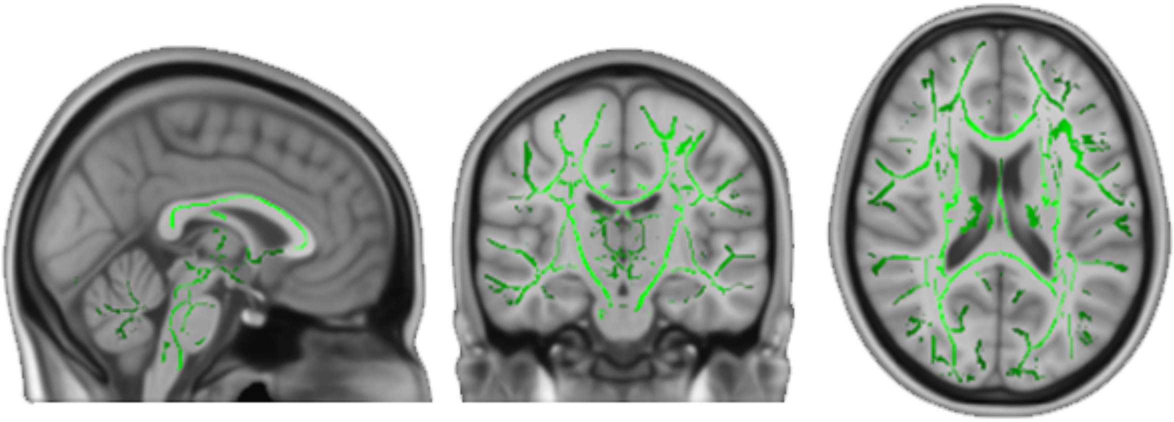

**Figure SM4. Results of the TBSS analysis**. TBSS results show no voxels of greater FA, MD, RD, or AD in either genotype group compared to the other. Displayed in green is the mean white matter skeleton for the analysed sample. The standard MNI152_1mm image is used as the background template.

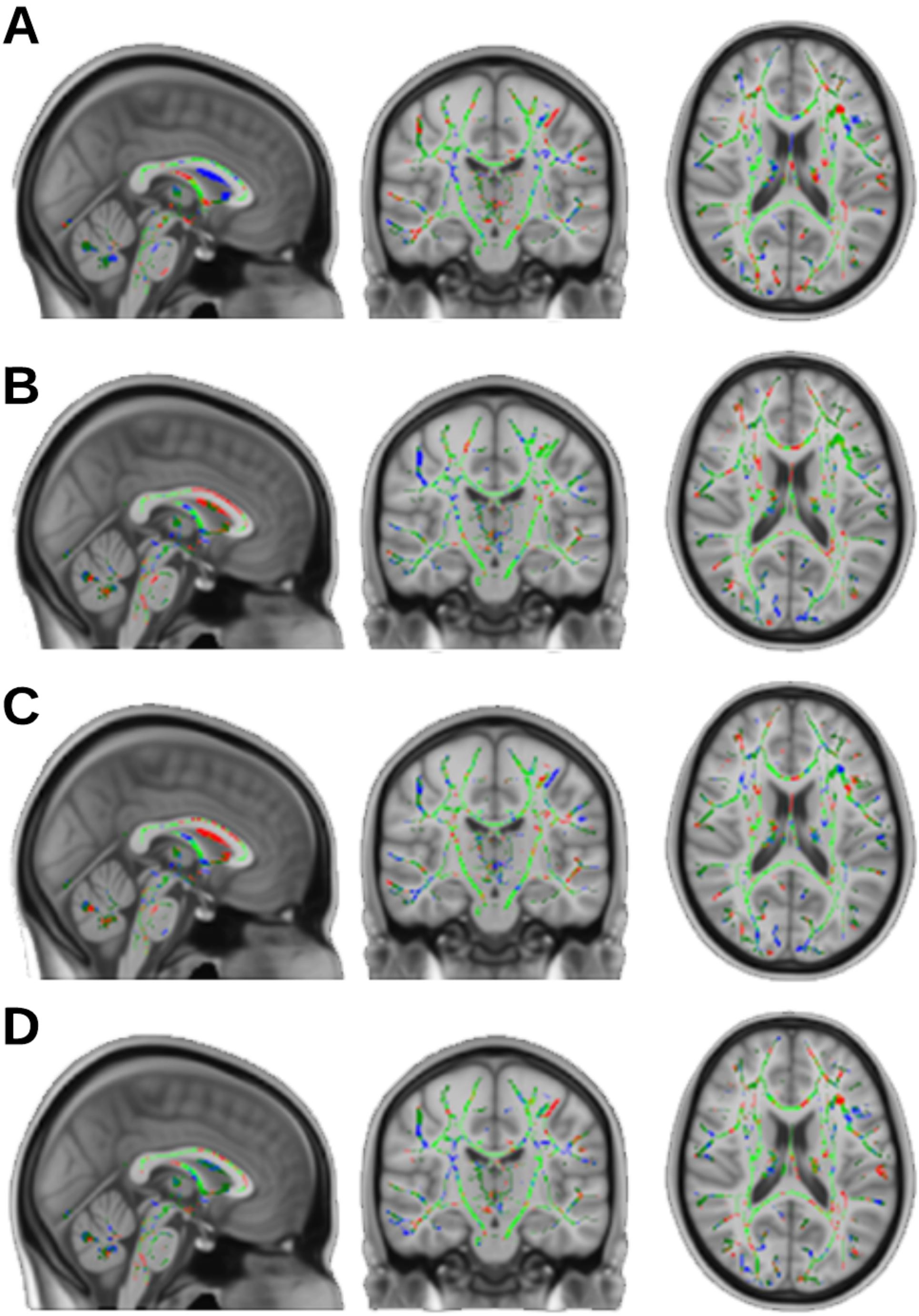


**Figure SM5. TBSS results prior to correction for multiple comparisons.** TBSS results prior to the corrections for multiple comparisons. Panel A: Fractional Anisotropy, Panel B:

Axial Diffusion, Panel C: Radial Diffusion, Panel D: Mean Diffusivity. Voxels in red are areas where Val/Val participants have greater values for the diffusivity parameter compared to Met+ carriers, while voxels in blue represent areas where Met+ carriers have greater values for the diffusivity parameter than Val/Val participants. Voxels showing no group differences are shown in green.

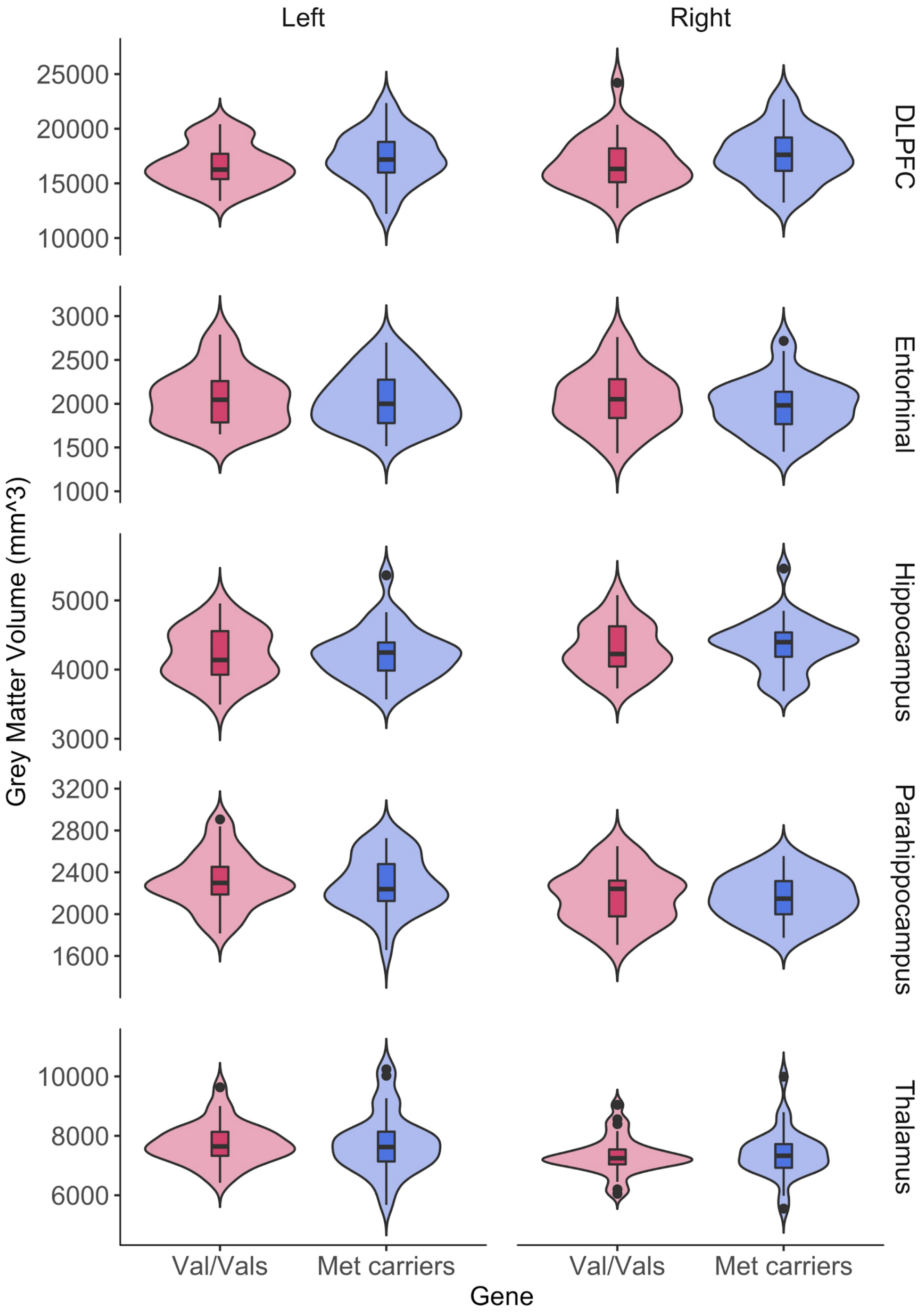


**Figure SM6. Grey matter volumes split by hemisphere.** Distributions of the grey matter volumes for each genotype (Val/Val vs Met+), across each structure (thalamus, parahippocampal region, hippocampus, entorhinal cortex, and dorsolateral prefrontal cortex), by hemisphere (left, right).

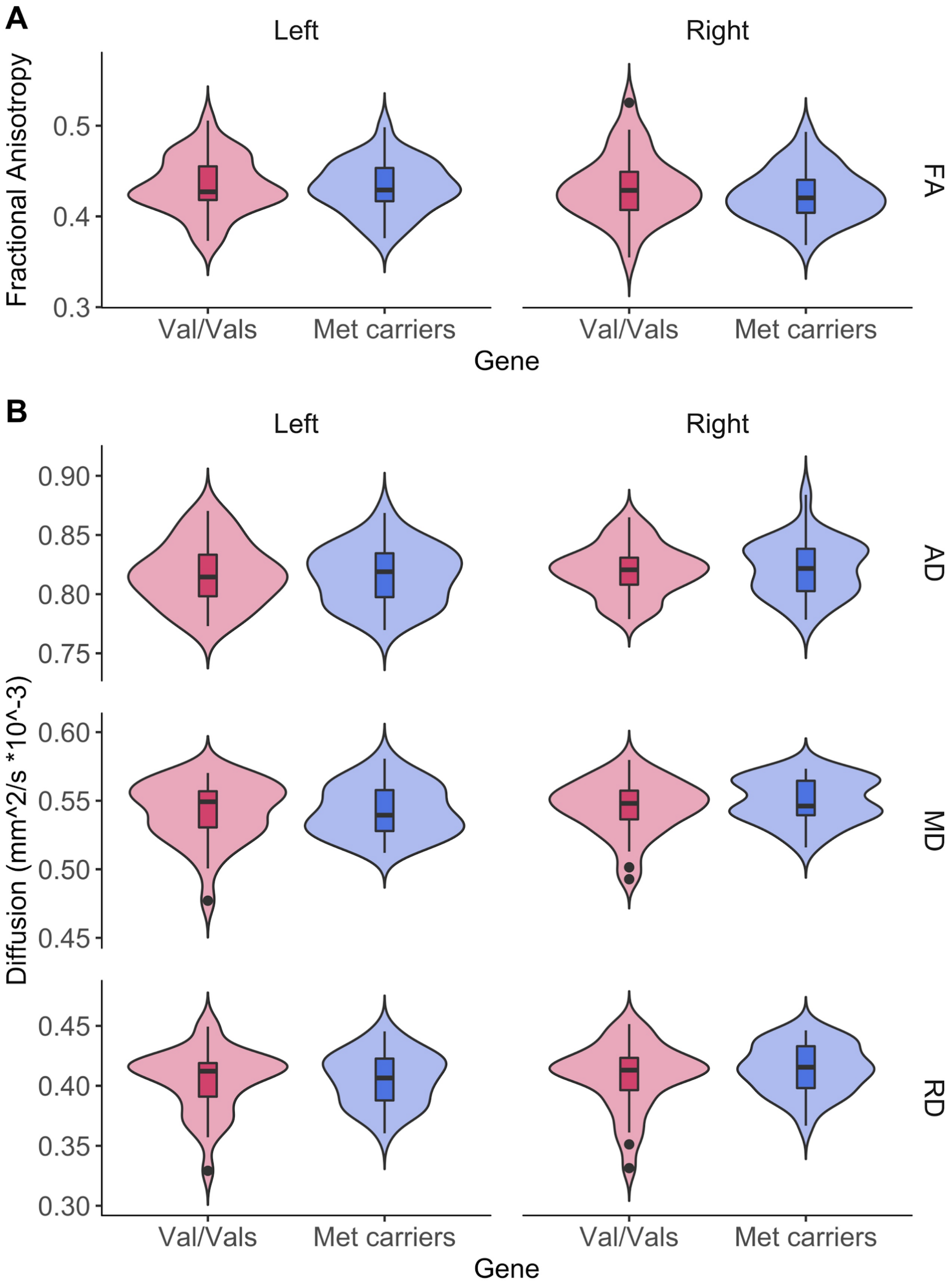


**Figure SM7. Uncinate fasciculus diffusion values split by hemisphere.** Distributions of the four diffusivity parameters (RD = radial diffusion, AD = axial diffusion, MD = mean diffusion, and FA = fractional anisotropy) measured across the uncinate fasciculus, by genotype, and hemisphere. In red are Val/Val participants, and in blue are Met allele carriers.

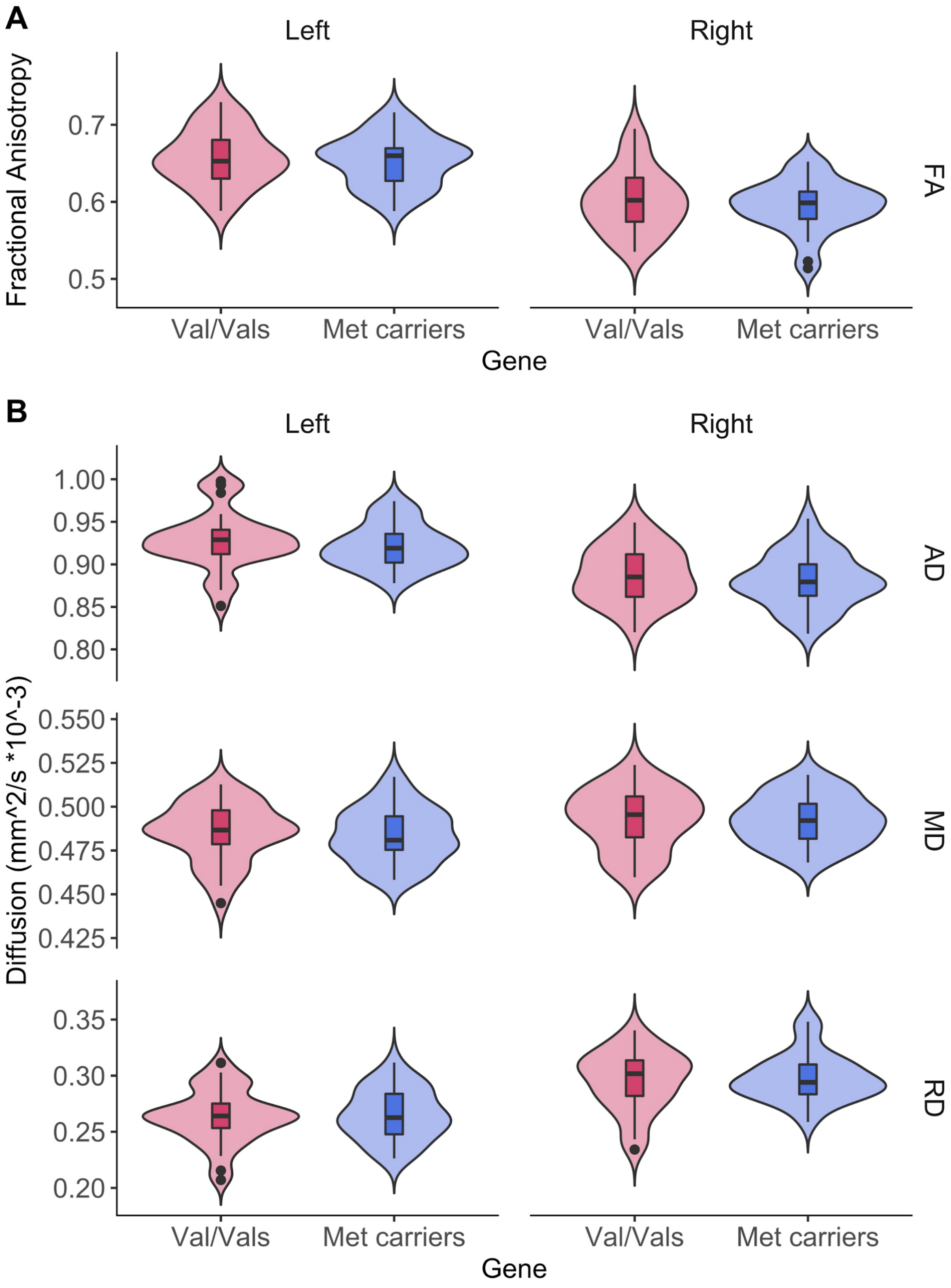


**Figure SM8. Cingulate gyrus diffusion values split by hemisphere.** Distributions of the four diffusivity parameters (RD = radial diffusion, AD = axial diffusion, MD = mean diffusion, and FA = fractional anisotropy) measured across the cingulate gyrus, by genotype, and hemisphere. In red are Val/Val participants, and in blue are Met allele carriers.

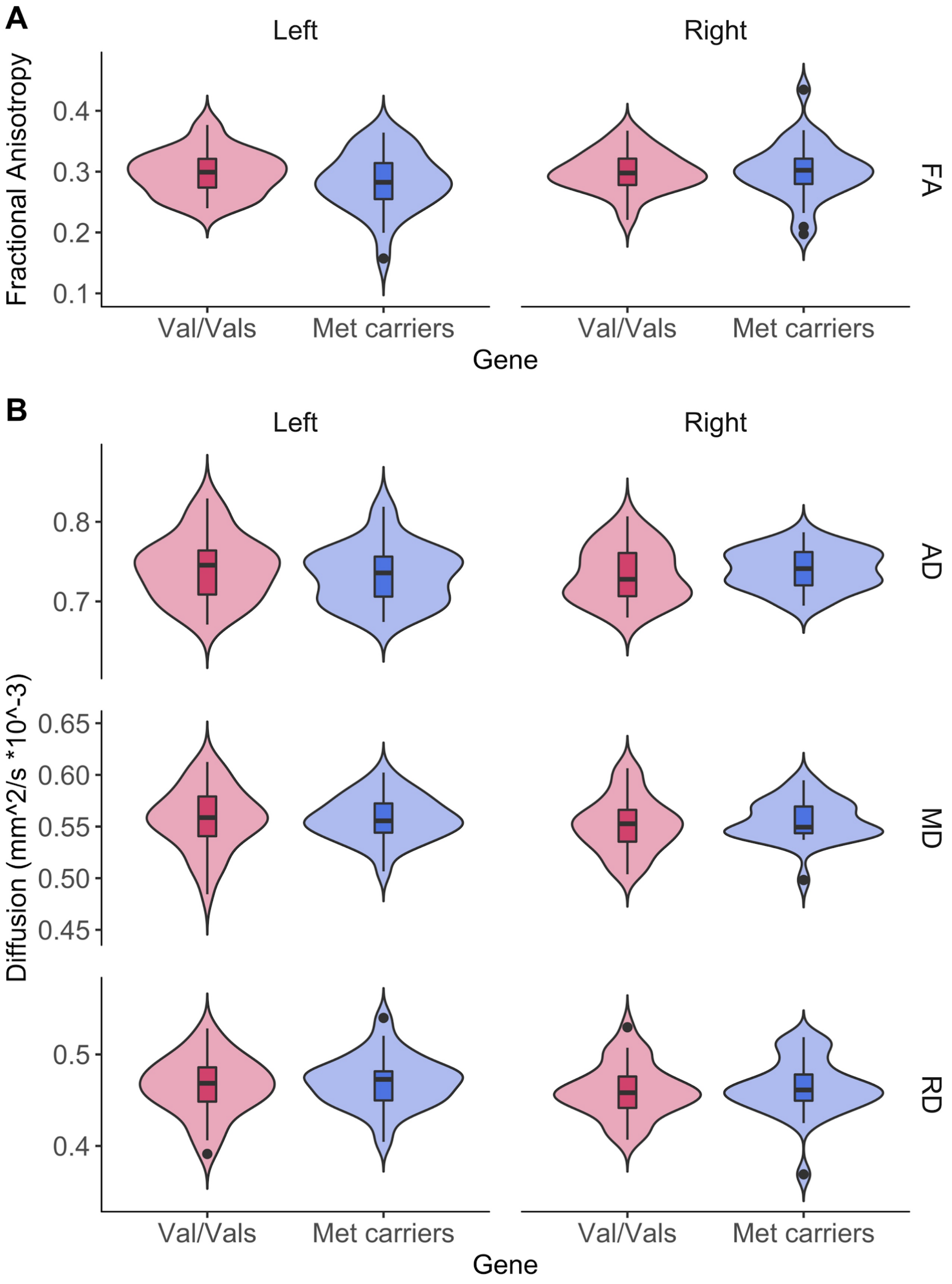


**Figure SM9. Cingulum angular bundle diffusion values split by hemisphere.**

Distributions of the four diffusivity parameters (RD = radial diffusion, AD = axial diffusion, MD = mean diffusion, and FA = fractional anisotropy) measured across the cingulum angular bundle, by genotype, and hemisphere. In red are Val/Val participants, and in blue are Met allele carriers.

#### Frequentist Alternative Analyses

Presented here are the frequentist equivalent analyses to the Bayesian analyses within the main body of text. In order to make these analyses as comparable as possible, we did not correct for multiple comparisons here.

**Behavioural analyses.** Two separate independent samples *t*-tests were used to compare the accuracy scores on our two tasks. For each test, we restricted the direction of testing, based on previous evidence that Val/Vals score higher on recognition tasks compared to Met allele carriers. On our measure of familiarity accuracy Val/Vals scored significantly higher than Met allele carriers ($t(57) = 1.95$, $p = 0.03$, $d = 0.51$, 95%CI = 0.07; ∞). However, for our recollection measure, there was no significant difference between the two groups ($t(57) = 0.79$, $p = 0.22$, $d = 0.21$, 95%CI = −0.22; ∞).

**Grey matter region of interest analyses.** A repeated measures ANOVA with mean grey matter volume values as a dependent variable, genotype (Val/Val, Met+) as the between subjects factor, structure (thalamus, parahippocampus, hippocampus, DLPFC, entorhinal cortex) as the within subjects factor, and eTIV and Age as covariates revealed no significant results. More specifically, there was no main effect of Val66Met genotype ($F(1, 57) = 0.01$, $p = 0.93$, $n^2_p < 0.01$). However, each of the covariates did significantly influence grey matter volume [eTIV: $F(1, 57) = 167.25$, $p < 0.01$, $n^2_p = 0.75$; Age: $F(1, 57) = 6.31$, $p = 0.02$, $n^2_p = 0.10$].

**White matter region of interest analyses.** Four separate repeated measures ANOVAs were conducted to look at how Val66Met genotype influences tract integrity, one for each diffusion parameter (FA, MD, RD, AD). A repeated measures ANOVA with mean FA values as a dependent variable, genotype (Val/Val, Met) as the between subjects factor, tract (uncinate fasciculus, cingulate gyrus, cingulum angular bundle) as the within subjects factor, and Age and estimated intracranial volume as covariates revealed no significant results of interest. More specifically, there was no main effect of the Val66Met genotype ($F(1, 57) = 0.58, p = 0.45, n^2_p < 0.01$) on FA values. However, each of the covariates did significantly influence grey matter volume [eTIV: $F(1, 57) = 4.81, p = 0.03, n^2_p = 0.08$; Age: $F(1, 57) = 1.56, p = 0.22, n^2_p = 0.03$]. Similar results are also observed for MD [Val66Met: $F(1, 57) = 0.09, p = 0.76, n^2_p < 0.01$; eTIV: $F(1, 57) = 8.53, p < 0.01, n^2_p = 0.13$; Age: $F(1, 57) = 15.60, p < 0.01, n^2_p = 0.24$], AD [Val66Met: $F(1, 57) = 1.25, p = 0.27, n^2_p = 0.02$; eTIV: $F(1, 57) = 0.96, p = 0.33, n^2_p = 0.02$; Age: $F(1, 57) = 9.74, p < 0.01, n^2_p = 0.15$], and RD [Val66Met: $F(1, 57) = 0.09, p < 0.76, n^2_p < 0.01$; eTIV: $F(1, 57) = 7.78, p < 0.01, n^2_p = 0.12$; Age: $F(1, 57) = 8.98, p < 0.01, n^2_p = 0.14$].

**Bayesian Alternative Analyses**

**Table SM1.** Bayes factors ($BF_{01}$) for the additional ANCOVAs run to include Sex and Ethnicity as covariates.

| Measure | ROI Tested | Sex | Ethnicity |
|---|---|---|---|
| GM Volume | Hippocampus | 3.67 | 2.61 |
| GM Volume | Parahippocampus | 3.40 | 3.20 |
| GM Volume | Entorhinal Cortex | 2.44 | 1.96 |
| GM Volume | Thalamus | 3.58 | 3.52 |
| GM Volume | DLPFC | 2.63 | 3.13 |
| FA | Uncinate Fasciculus | 2.46 | 3.49 |
| FA | Cingulate Gyrus | 3.20 | 3.48 |
| FA | Cingulum Angular Bundle | 2.95 | 3.46 |
| MD | Uncinate Fasciculus | 3.56 | 3.70 |
| MD | Cingulate Gyrus | 3.02 | 2.17 |
| MD | Cingulum Angular Bundle | 3.66 | 3.55 |
| AD | Uncinate Fasciculus | 3.07 | 3.55 |
| AD | Cingulate Gyrus | 1.76 | 3.71 |
| AD | Cingulum Angular Bundle | 3.22 | 3.66 |
| RD | Uncinate Fasciculus | 3.09 | 3.61 |
| RD | Cingulate Gyrus | 3.61 | 3.77 |
| RD | Cingulum Angular Bundle | 3.59 | 3.59 |
